# Notch Regulates Vascular Collagen IV Basement Membrane Through Modulation of Lysyl Hydroxylase 3 Trafficking

**DOI:** 10.1101/2020.06.22.165431

**Authors:** Stephen J. Gross, Amelia M. Webb, Alek D. Peterlin, Jessica R. Durrant, Rachel Judson, Erich J. Kushner

**Author notes:** Author for correspondence: Erich Kushner, University of Denver, Department of Biological Sciences, Denver, CO 80210.

## Abstract

During angiogenesis, endothelial cells secrete proteins that make up a planar protein network surrounding blood vessels termed basement membrane (BM). Collagen type IV (Col IV) is a BM protein associated with early blood vessel morphogenesis and is essential for blood vessel stability. To date, little is known about how endothelial cells mediate intracellular transport and selective secretion of Col IV. We have identified the GTPase Rab10 as a major regulator of Col IV vesicular trafficking during vascular development. Knockdown of Rab10 reduced *de novo* Col IV secretion *in vivo* and *in vitro*. Mechanistically, we determined that Rab10 is an indirect mediator of Col IV secretion, partnering with atypical Rab25 to deliver the enzyme lysyl hydroxylase 3 (LH3) to Col IV-containing vesicles staged for secretion. Loss of Rab10 or Rab25 resulted in depletion of LH3 from Col IV-containing vesicles and rapid lysosomal degradation of Col IV. Furthermore, we demonstrated that Rab10 activation is downstream of Notch signaling, indicating a novel connection between permissive Notch-based vessel maturation programs and vesicle trafficking. Overall, our results illustrate both a new trafficking-based component in the regulated secretion of Col IV and how this vesicle trafficking program interfaces with Notch signaling to fine-tune BM secretion during blood vessel development.

## INTRODUCTION

Endothelial cells (ECs) are the cell type responsible for the bulk of embryonic blood vessel formation, eventually leading to an estimated 50,000 miles of vasculature by adulthood[1]. During development, new blood vessels emerge from pre-existing vasculature, a process termed angiogenesis [1-3]. During angiogenesis, ECs secrete a variety of proteins composing a planar protein network that encapsulates blood vessels, collectively termed basement membrane (BM). The bulk of the vascular BM is secreted during the angiogenic stages of development by ECs and later buttressed with mural cell interactions[4]. The BM not only provides a 50-200 nm thick static planar protein network on which ECs reside, but constitutes a dynamic and diverse extracellular environment vital to blood vessel integrity[5, 6]. The perivascular BM elements vary depending on anatomical location[7], but generally demonstrate an enrichment of macromolecular collagen IV (Col IV), laminins (4-1-1 and 5-1-1), perlecan, fibronectin and nidogen[4, 8] that are directly secreted by ECs[9] and supportive cells [9-11]. For instance, laminins are anchored to Col IV by cross-linking of perlecan and nidogen, creating an exceptionally resilient co-polymer[12, 13]. In the absence of Col IV, BM integrity and cell integrin signaling are greatly diminished[14, 15]. In sprouting angiogenesis, ECs break down existing BM while simultaneously secreting it. How ECs orchestrate this feat, blending cell-autonomous signaling with tissue-level communication, is unclear and represents a void in our understanding of blood vessel development.

Blood vessels are exquisitely dependent on Col IV BM due their inherent pressure demands as a fluid transport system. Disruption in Col IV bioavailability during blood vessel development is the basis of small vessel disease (SVD) in which Col IV point mutations promote intracellular retention or degradation of Col IV, limiting its perivascular deposition. This reduced Col IV secretion in SVD is associated with a clinical sequela like intracerebral hemorrhage, typically resulting in death or profound disability[16]. Genetic ablation of Col IV in mice does not prevent angiogenesis, *per se*, but is embryonically lethal due to an inability to resist the mechanical strain of blood circulation and resulting vessel rupture[15]. Col IV itself is an obligate heterotrimer made of 3 alpha chains forming a long triple helix[17]. Trimer formation is, in part, achieved through lysyl hydroxylase (LH) 1-3 (genes Procollagen-Lysine, 2-Oxoglutarate 5-Dioxygenase (PLOD1,2,3)) that catalyze hydroxylysine formation, without which stable heterotrimer formation is abolished. LH1 and LH2 are restricted to the endoplasmic reticulum (ER). LH3 demonstrates an affinity for Col IV over other collagen subtypes and is found both in the ER and on post-Golgi vesicles[18]. Indeed, Col IV homeostasis is important for both blood vessel development and maintenance.

Akin to transcriptional networks, vesicular trafficking programs are complex and likely comprised of unique organotypic signatures that are fundamental to tissue form and function. In terms of Col IV, how Col IV is transported, targeted to the basal membrane, interfaces with degradative organelles or intersects with other proteins/enzymes during angiogenesis is mostly unknown. Moreover, it is well characterized that permissive programs, such as Notch signaling[19] are indispensable for blood vessel maturation and stability. Taken in aggregate, how blood vessel maturation signaling initiates crosstalk with trafficking regulators, such as those involved in Col IV secretion, is also a major void in our understanding of BM regulation during angiogenesis.

Here, we describe a novel level of Col IV regulation that leverages vesicular transport to precisely modulate Col IV secretion in ECs during blood vessel development. Specifically, we demonstrate that Rab10 and Rab25 GTPases govern the transport and fusion of LH3 to Col IV vesicles staged for secretion. In the absence or inactivation of Rab10 or Rab25, LH3 trafficking is halted and Col IV secretion is abolished. Additionally, we demonstrate a first of its kind connection between permissive Notch signaling and control of LH3 trafficking to regulate Col IV bioavailability during blood vessel development. We demonstrate that a Rab guanine exchange factor (GEF) is a downstream transcriptional target of Notch activation, linking Notch signaling to vesicular trafficking regulation of Col IV during blood vessel development.

## RESULTS

### Loss of Rab10 impairs endothelial basement membrane secretion

Based on previous literature implicating the GTPase Rab10 as a Col IV trafficking mediator, we first sought to determine if Rab10 was involved in Col IV secretion in primary ECs[5]. ECs plated on coverslips demonstrated a robust secretion of Col IV marked by long trails leading back to individual ECs. Knockdown of Rab10 significantly diminished Col IV secretion with a limited amount of Col IV being deposited under the ventral/basal surface of the EC (**Figure 1A-1C**). Next, we transduced ECs with wild-type (WT), constitutively active (CA, Q68L) or a dominant negative (DN, T23N) Rab10 fused to a green fluorescence protein (GFP). ECs expressing a GFP-Rab10 WT or CA mutant did not show any difference in Col IV secretion compared with each other. However, expression of the GFP-Rab10 DN significantly reduced Col IV secretion compared to both WT and CA Rab10 (**Figure 1D and 1E**). Secreted Col IV is typically associated with other BM proteins such as perlecan and laminin[20, 21]. Knockdown of Rab10 also blunted the secretion of these vascular BM proteins (**Figure 1F and 1G**). It is possible that reduced secretion could be related to diminished migratory capacity in ECs lacking Rab10. To factor this out, we performed a scratch wound assay as a gauge of cell motility. There was no effect of Rab10 knockdown on cell migration (**Figure S1A and S1B**). Additionally, we determined that Rab10 did not affect apoptotic tendency by measuring cleaved caspase-3 levels **(Figure S1C and S1D**). These results suggest that Rab10 is associated with Col IV secretion in ECs.

**Figure 1.**
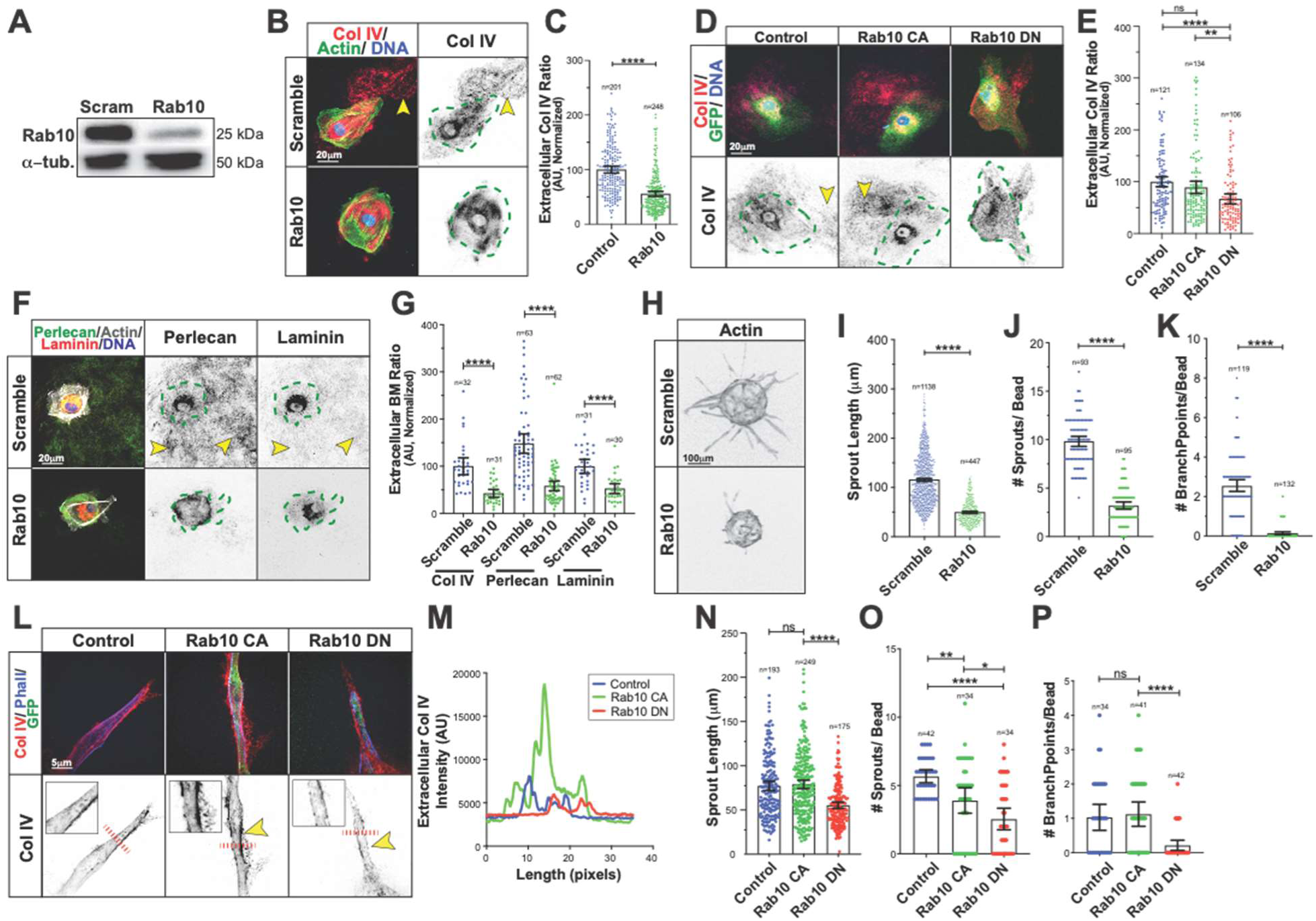
Loss of Rab10 impairs endothelial basement membrane secretion. (A) Immunoblot of Rab10 in ECs transfected with either scramble or Rab10 siRNA and probed for indicated proteins. (B) Representative images of scramble or Rab10 siRNA-treated ECs and stained for collagen IV (Col IV) (red), actin (green), and DNA (blue). Dotted green line indicates cell outline. Arrowheads denote extracellular Col IV secretion. (C) Graph of extracellular Col IV ratio in scramble or Rab10 siRNA-treated ECs. n = number of ECs. (D) Representative image of ECs expressing GFP only (control), GFP-Rab10 constitutively active (CA) or GFP-Rab10 dominant negative (DN), stained for collagen IV (red) and DNA (blue). Dotted green line indicates cell outline. Arrowheads denote extracellular Col IV secretion. (E) Graph of extracellular Col IV ratio in ECs expressing GFP only (control) or GFP Rab10 CA/DN. n=number of ECs. (F) Representative images of scramble or Rab10 siRNA-treated ECs and stained for laminin (red) perlecan (green), actin (grey), and DNA (blue). Dotted green line indicates cell outline. Arrowheads denote extracellular Col IV secretion. (G) Graph of extracellular basement membrane (BM) ratio in scramble or Rab10 siRNA-treated ECs between indicated secreted proteins. n=number of ECs. (H) Representative image of fibrin-bead sprouts between indicated siRNA treatment groups. Sprouts were stained for actin (grey) to delineate sprout morphology. (J-K) Graphs of sprouting parameters for scramble or Rab10 siRNA-treated sprouts. n=number of measurements. (L) Representative images of ECs expressing GFP only (control), GFP-Rab10 CA or GFP-Rab10 DN in fibrin-bead sprouts. Dotted red line indicates line scan of (m). Arrowheads denote extracellular Col IV secretion. (M) Line scan measurement of Col IV fluorescence intensity shown in (L), dotted red lines. (N-P) Graphs of sprouting parameters for GFP only (control) or GFP-Rab10 CA/DN expressing sprouts. n=number of measurements. For all experiments, data are represented as mean ± 95% confidence intervals. Black bars indicate comparison groups with indicated p-values. All p-values are from two-tailed Student’s t-test from at least three experiments. *p≤0.05; **p≤0.01; ***p≤0.001; ****p≤0.0001; ns, not significant.

To determine if Rab10 affected 3-dimensional (3D) sprouting behaviors we employed a fibrin-bead assay in which ECs form multicellular sprouts in a fibrin matrix[22]. Loss of Rab10 posed a severe detriment on sprouting behaviors with a 70-90% reduction in sprout parameters compared with controls (**Figure 1H-1K**). To further probe how loss and gain of function of Rab10 affected Col IV secretion in 3D sprouting, we mosaically transduced GFP-Rab10 DN and CA mutants into growing sprouts. Staining non-permeabilized sprouts for secreted Col IV showed that ECs expressing the DN form of Rab10 had lower levels of perivascular Col IV, while the CA Rab10 mutant showed qualitatively elevated Col IV secretion compared with a non-transduced control (**Figure 1L and 1M**). In comparing sprout morphology, only the DN version of Rab10 impaired sprout length and number of branch points in reference to WT and CA Rab10 expressing ECs (**Figure 1N and 1P)**. Interestingly, both Rab10 CA and DN impaired sprout formation, the DN variant to a greater magnitude, compared with WT Rab10 (**Figure 1O**). These results indicate that Rab10 is necessary for *in vitro* sprouting. Additionally, these data suggest that the relative level of Rab10 activation may also be consequential for proper sprout formation.

### Rab10 influences Col IV bioavailability *in vivo*

To explore if Rab10 was required *in vivo* we turned to a mouse model of blood vessel development (**Figure S2A**). Homozygous loss of Rab10 was lethal at embryonic day 7, consistent with other reports[23]. Rab10 heterozygous mice (Rab10^em1(IMPC)J^) were viable and did not show any appreciable differences in survival compared with WT littermates. To examine if Rab10 heterozygosity impacted Col IV bioavailability and, subsequently, sprouting angiogenesis we examined the retinal vascular plexus[24]. There was no difference in sprouting parameters between Rab10^+/-^ and WT littermates at postnatal (P) day 6 (**Figure 2A-C**). However, there was a ∼18% reduction in vascular Col IV intensity in the Rab10^+/-^ group compared with WTs, indicating Col IV secretion was slightly reduced with only one working Rab10 allele (**Figure 2D**). We confirmed this reduced Col IV abundance in serial sections of intracranial blood vessels alternating between H&E staining and Col IV immunohistochemistry to compare the same anatomical location between groups (**Figure 2E**). Other vessel beds, such as those in the dermal tissue showed a similar reduction in Col IV staining (**Figure S2B**). A fraction of specimens collected at P6 exhibited cerebral hemorrhage (**Figure S2C**) potentially suggesting compromised blood vessel integrity; although, this phenotype was not highly penetrant. These data suggest that Rab10 haploinsufficiency is associated with reduced perivascular Col IV.

**Figure 2.**
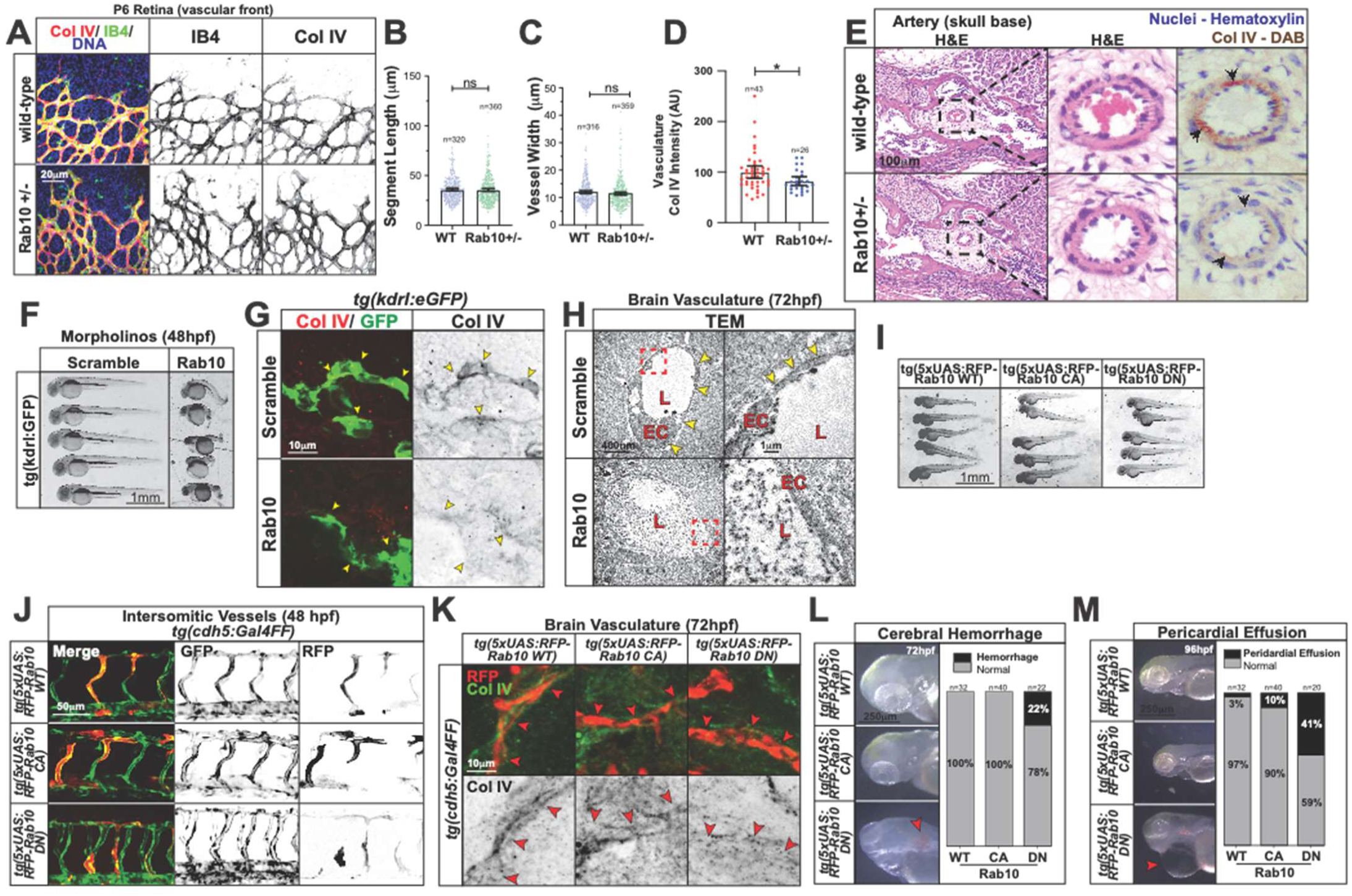
Rab10 influences Col IV bioavailability *in vivo*. (A) Representative images of WT or Rab10^+/-^ mouse retinas harvested at P6. Retinas were stained for Col IV (red), DNA (blue), and conjugated-isolectin B4 (IB4) (green) to identify blood vessels. (B,C) Graph of sprouting parameters for WT or Rab10^+/-^ P6 mouse retinas. (D) Graph of vasculature Col IV fluorescence intensity for WT or Rab10^+/-^ P6 mouse retinas. (E) Hematoxylin and eosin (H&E) stained tissue slices from WT or Rab10^+/-^ P6 mice. Col IV stained with DAB (3-3’ diaminobenzidine) indicated by arrowheads. (F) Representative images of 48 hpf *tg(kdrl:eGFP)* zebrafish injected with scramble or Rab10 morpholinos. (G) Representative images from brain cryosections of 48 hpf *tg(kdrl:eGFP)* zebrafish from scramble or Rab10 morphants. Arrowheads denote Col IV extracellular secretion. (H) Transmission electron microscopy of 72 hpf *tg(kdrl:eGFP)* zebrafish from scramble or Rab10 morphants. Dotted red box indicates enlargement window. Arrowheads denote basement membrane deposition. EC, endothelial cell; L, lumen. (I) Representative images of 48 hpf *tg(cdh5:gal4ff*) zebrafish expressing 5xUAS:RFP-Rab10 WT, CA or DN. (J) Representative images of 48 hpf *tg(cdh5:gal4ff*) zebrafish expressing 5xUAS:RFP-Rab10 WT, CA or DN in intersomitic blood vessels. (K) Representative images of 72 hpf *tg(cdh5:gal4ff*) zebrafish expressing 5xUAS:RFP-Rab10 WT/CA/DN brain cryosections. (L) Representative images of 72 hpf *tg(cdh5:gal4ff*) zebrafish with 5xUAS:RFP-Rab10 WT/CA/DN injections. Arrowhead denotes cerebral hemorrhage. (M) Representative images of 96 hpf *tg(cdh5:gal4ff*) zebrafish with 5xUAS:RFP-Rab10 WT/CA/DN injections. Arrowhead denotes pericardial effusion. For all experiments, data are represented as mean ± 95% confidence intervals. Black bars indicate comparison groups with indicated p-values. All p-values are from two-tailed Student’s t-test from at least three experiments. *p≤0.05; **p≤0.01; ***p≤0.001; ****p≤0.0001; ns, not significant.

Given Rab10 null mice were not viable, we moved to a zebrafish model to more easily gain access to earlier stages of development. The zebrafish Rab10 ortholog is 97% identical to the human ortholog (**Figure S2D**). Morpholino knockdown of Rab10 resulted in developmental defects, primarily a severe dorsalized phenotype compared with scrambled controls (**Figure 2F**). Sectioning of morphant embryos expressing a vascular reporter *tg(kdrl:eGFP)* with normal cranial development at 72 hours post fertilization (hpf) showed a marked reduction in perivascular Col IV levels (**Figure 2G**). Confirming this observation, using transmission electron microscopy we observed that Rab10 morphants exhibited little to no basal lamina surrounding brain ECs compared with controls (**Figure 2H**). To subvert the effect of global Rab10 loss of function, we mosaically over-expressed TagRFP-tagged Rab10 WT, CA or DN in the blood vessels of 72 hpf zebrafish. The resulting zebrafish showed mosaic vascular expression of Rab10 variants with no visible impact on body plan or the intersomitic vessels (**Figure 2I and 2J**). Again, staining for Col IV in the brain vasculature, we observed that the Rab10 DN mutant alone reduced perivascular Col IV compared with overexpression of WT and CA Rab10 constructs (**Figure 2K**). Strikingly, at 72 hpf fish injected with the DN Rab10 demonstrated elevated frequencies of cerebral hemorrhage and pericardial effusion, suggesting compromised blood vessel integrity (**Fig. 2L and 2M**). These results indicate that loss of Rab10 impairs Col IV bioavailability.

### Rab10 influences intracellular Col IV protein stability

Next, we sought to understand how Rab10 impacts Col IV secretion by first investigating normal Col IV cellular turnover in ECs. ECs incubated with the Golgi-disrupting compound brefeldin A (BFA) ablated Col IV secretion, suggesting that Col IV itself or its regulators use a classical post-Golgi trafficking route (**Figure 3A and 3B**). Next, cycloheximide (CHX) with and without BFA was added to inhibit new protein synthesis to determine the half-life of the intracellular Col IV pool in the absence of secretion. Cycloheximide addition reduced the intracellular Col IV pool by 75% at 4 hours (**Figure 3C**). Strikingly, addition of the secretion inhibitor BFA doubled the Col IV content in the cycloheximide condition (**Figure 3D and 3E**), indicating that at least half of the Col IV decay was due to secretion. Knockdown of Rab10 closely mimicked BFA-induced intracellular Col IV retention (**Figure 3F and 3G**). These results demonstrate the EC Col IV pool is rapidly secreted and knockdown of Rab10 resembles chemically induced inhibition of secretion, thus Rab10 may play a mechanistic role in this pathway.

**Figure 3.**
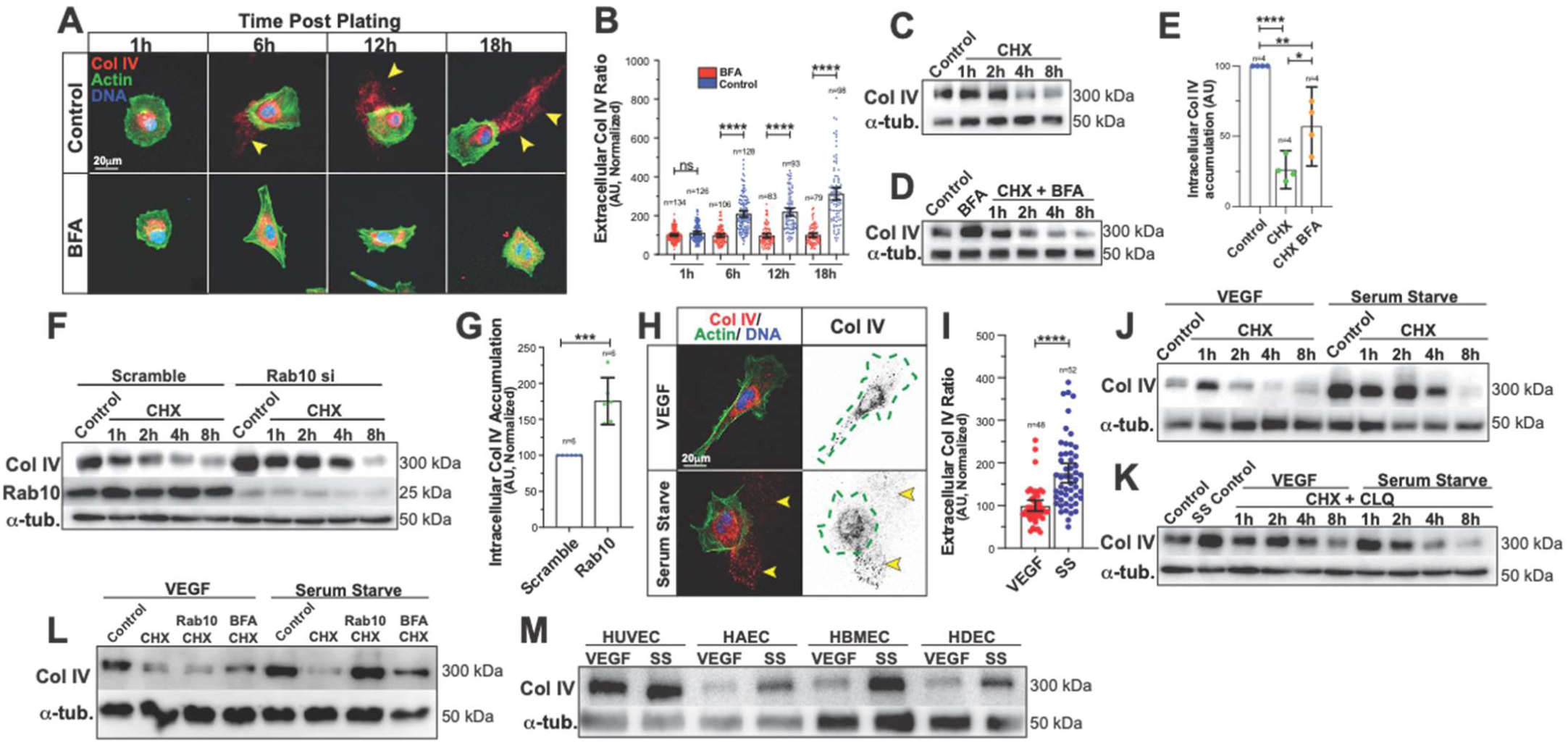
Rab10 influences intracellular Col IV protein stability. (A) Representative images of ECs treated with brefeldin A (BFA) and fixed at indicated time points after BFA exposure. Cells were stained for Col IV (red), actin (green), and DNA (blue). Arrowheads denote extracellular Col IV secretion. (B) Graph of extracellular Col IV ratio in control or BFA-treated ECs at indicated time points after BFA exposure. (C) Immunoblot of Col IV in ECs treated with cycloheximide (CHX) (20mg/ml) and probed for indicated proteins. (D) Immunoblot of Col IV in ECs treated with both CHX and BFA. (E) Graph of Col IV protein accumulation in ECs treated with CHX, and CHX with BFA. Col IV accumulations were normalized to α-tubulin levels. (F) Immunoblot of Col IV in ECs transfected with either scramble or Rab10 siRNA and treated with CHX. EC lysate probed for indicated proteins. (G) Graph of Col IV levels in ECs transfected with either scramble or Rab10 siRNA and treated with CHX. Col IV accumulations were normalized to α-tubulin levels. (H) Representative images of ECs cultured in VEGF-containing or serum starve (SS) media and stained for collagen IV (red), actin (green), and DNA (blue). Dotted green line indicates cell outline. Arrowheads denote extracellular Col IV secretion. (I) Graph of extracellular Col IV ratio of ECs cultured in VEGF-containing or SS media. (J) Immunoblot of Col IV in ECs cultured in VEGF-containing or SS media treated with CHX at indicated time points and probed for indicated proteins. (K) Immunoblot of Col IV in ECs cultured in VEGF-containing or SS media treated with CHX and chloroquine (CLQ) (10µM) at indicated time points and probed for indicated proteins. (L) Immunoblot of Col IV in ECs transfected with either scramble or Rab10 siRNA and cultured in VEGF-containing or SS media treated with CHX and/or BFA for 8 hrs and probed for indicated proteins. (M) Immunoblot of Col IV in various ECs (HUVECs, HAECs, HBMECs, and HDECs) cultured in either VEGF-containing or SS media and probed for indicated proteins. For all experiments, data are represented as mean ± 95% confidence intervals. Black bars indicate comparison groups with indicated p-values. All p-values are from two-tailed Student’s t-test from at least three experiments. *p≤0.05; **p≤0.01; ***p≤0.001; ****p≤0.0001; ns, not significant.

Vascular endothelial growth factor (VEGF) is one of the most well-characterized proangiogenic factors and is essential for angiogenesis[24-28]. Given VEGF signaling is required for embryonic blood vessel development, we sought to determine how VEGF influenced Col IV secretion. Strikingly, VEGF ligand administration significantly impeded Col IV secretion in freely migrating ECs compared with ECs in serum-starvation (SS) culture media (**Figure 3H and 3I**). Moreover, VEGF supplementation showed a concentration-dependent reduction in Col IV, laminin and perlecan secretion (**Figure S3**). Once again, we used cycloheximide to determine the half-life of intracellular Col IV with VEGF stimulation. In line with the general lack of secretion in the VEGF-treated ECs, VEGF stimulation depleted the Col IV intracellular pool compared with a SS control (**Figure 3J**). Incubation with the lysosomal inhibitor chloroquine (CLQ) rescued VEGF-mediated loss of Col IV, indicating that VEGF induced Col IV destruction via the lysosome **(Figure 3K**). We next compared how Rab10 affected intracellular Col IV levels in VEGF-exposed and SS states. Knockdown of Rab10 in the presence of VEGF still resulted in Col IV degradation as compared with the SS state or BFA control when treated with cycloheximide (**Figure 3L**). This finding promotes the notion that VEGF-induced Col IV degradation does not involve Rab10, suggesting that Rab10 is participating in a secretory, not a degradative pathway. To ensure this degradative response to VEGF was not restricted to human umbilical vein ECs, we assayed for intracellular Col IV levels in human aortic, human brain, human microvascular and human dermal primary ECs. Across all primary cell lines, Col IV levels were reduced in the VEGF-treated state compared with SS ECs (**Figure 3M**), suggesting this is likely a global endothelial response.

### Rab10 and Rab25 work in combination to traffic LH3 to collagen IV containing vesicles

Given the strong effect of Rab10 on both Col IV secretion and bioavailability, we originally hypothesized that Rab10 was directly mediating Col IV vesicular trafficking (e.g. directly attached to Col IV vesicles). However, we did not observe Rab10 co-localization with Col IV-containing (CIVC) vesicles (**Figure 4A**). This lack of co-localization elevated the hypothesis that Rab10 may play an indirect role in Col IV trafficking. To this end, the lysyl hydroxylase 3 (LH3) enzyme has been shown to be critical for Col IV secretion and post-Golgi protein stability[29]. Additionally, it has been previously reported that LH3 trafficking requires both Rab10 and Rab25[30]. To determine if Rab10 was involved in the LH3 trafficking itinerary, we first compared Col IV secretion between Rab10 and Rab25 knockdowns to LH3 knockdowns. Loss of Rab10 or Rab25 phenocopied the reduced EC Col IV secretion observed with LH3 KD (**Figure 4B and 4C**), indicating that both Rab10 and Rab25 impact Col IV secretion to a similar magnitude compared with LH3 depletion. Rab10 and Rab25 depletion also significantly reduced sprouting parameters similar to LH3 knockdown in reference to a control group (**Figure 4D-4G**). These data demonstrate that Rab10 and Rab25 equally affect Col IV secretion and sprouting parameters in comparison to LH3 knockdown.

**Figure 4.**
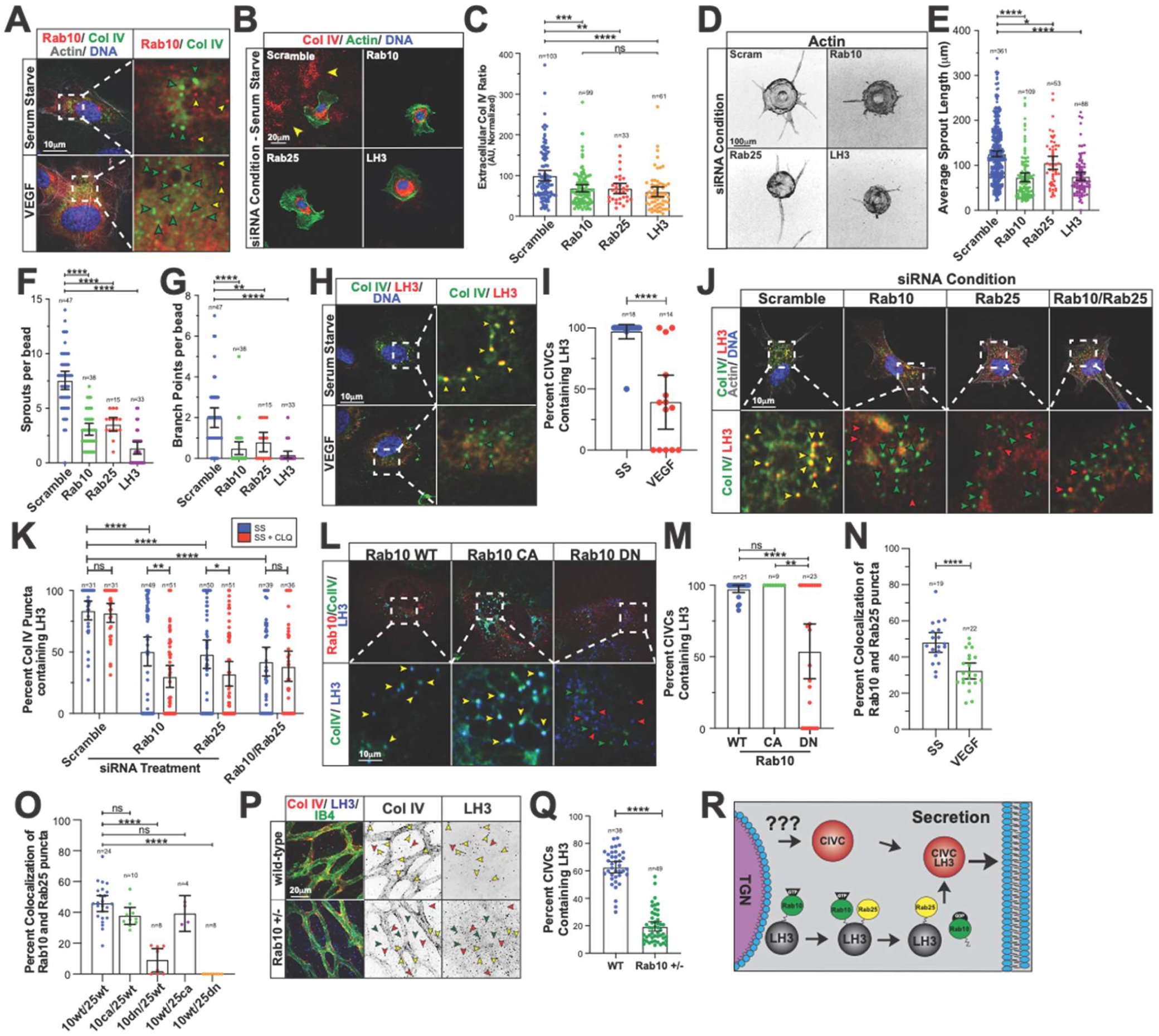
Rab10 and Rab25 work in combination to traffic LH3 to CIVCs. (A) Representative images of ECs expressing RFP-Rab10 WT and stained for Col IV (green). Green arrowheads indicate CIVCs only and yellow arrowheads indicate Rab10 puncta only. (B) Representative images of scramble, Rab10, Rab25, or LH3 siRNA-treated ECs and stained for Col IV (red), actin (green), and DNA (blue). Arrowheads denote extracellular Col IV secretion. (C) Graph of extracellular Col IV ratio of scramble, Rab10, Rab25, or LH3 siRNA-treated ECs cultured in SS media. (D) Representative images of fibrin-bead sprouts between indicated siRNA treatment groups. Sprouts were stained for actin (grey) to delineate sprout morphology. (E-G) Graphs of sprouting parameters for scramble, Rab10, Rab25, or LH3 siRNA-treated sprouts. (H) Representative images of ECs cultured in VEGF-containing or SS media and stained for Col IV (green), LH3 (red), and DNA (blue). Green arrowheads indicate CIVCs only and yellow arrowheads indicate co-localized puncta. (I) Graph of percent CIVC vesicles co-localized with LH3 in ECs cultured in VEGF-containing or SS media. (J) Representative images of scramble, Rab10, Rab25, or both Rab10/25 siRNA-treated ECs and stained for Col IV (green), LH3 (red), actin (grey), and DNA (blue). Yellow arrowheads indicate co-localized puncta only, green arrowheads indicate Col IV only puncta, and red arrowheads indicate LH3 puncta only. (K) Graph of percent CIVC vesicles co-localized with LH3 in scramble, Rab10, Rab25, or LH3 siRNA-treated conditions cultured in SS media or SS media with CLQ (10µM). (L) Representative images of ECs expressing RFP-Rab10 WT, CA, DN, stained for Col IV (green) and LH3 (blue). Yellow arrowheads indicate co-localized puncta only, green arrowheads indicate Col IV puncta, and red arrowheads indicate LH3 puncta. (M) Graph of percent CIVC vesicles co-localized with LH3 in ECs expressing RFP-Rab10 WT, CA or DN. (N) Graph of percent Rab10 puncta co-localized with Rab25 in either VEGF containing or SS media. (O) Graph of percent Rab10 puncta co-localized with Rab25 in ECs transfected with indicated constructs. (P) Representative images of WT or Rab10^+/-^ mice retinas harvested at P6. Retinas were stained for Col IV (red), LH3 (blue) and IB4 (green) to identify blood vessels. (Q) Graph of percent CIVCs containing LH3 in WT or Rab10^+/-^ P6 mouse retinas. (R) Schematic diagram showing how Rab10 and Rab25 function to coordinate delivery of LH3 to CIVCs for proper secretion of Col IV. Trans-Golgi network (TGN). For all experiments, data are represented as mean ± 95% confidence intervals. Black bars indicate comparison groups with indicated p-values. All p-values are from two-tailed Student’s t-test from at least three experiments. *p≤0.05; **p≤0.01; ***p≤0.001; ****p≤0.0001; ns, not significant.

Reasoning that LH3 is the cargo of Rab10 and Rab25, we first determined the efficiency of LH3 transport to CIVC vesicles in the VEGF-treated and SS state in which Col IV secretion is greatly affected. We observed that in the SS state, LH3 co-localized with CIVC vesicles 99% of the time, while VEGF-treatment reduced LH3/CIVC vesicle co-localization by ∼60% (**Figure 4H and 4I**). Given the SS culture condition produced a near perfect co-localization between LH3 and CIVC vesicles, we used this SS condition to test how loss of Rab10 and Rab25 trafficking impacts LH3 transport to CIVC vesicles. Strikingly, knockdown of Rab10, Rab25 or combination significantly reduced LH3 and CIVC vesicle co-localization compared with controls (**Figure 4J and 4K**). In this experiment, chloroquine was added to prevent both LH3 and Col IV degradation to determine what fraction may be lysosomally degraded when Rab10 and Rab25 trafficking mediators are absent. Interestingly, lysosome inhibition significantly reduced the percentage of co-localization of LH3 and CIVC vesicles in Rab10 and Rab25, but not in double knockdown groups suggesting that a fraction of the LH3 or Col IV pool is degraded when trafficking is disrupted (**Figure 4K**). We next expressed WT, CA and DN Rab10 versions in ECs cultured in SS media. Wild-type and CA Rab10 overexpression did not affect LH3 transport to CIVC vesicles; however, the DN Rab10 mutant alone reduced LH3/CIVC vesicle co-localization by 50% (**Figure 4L and 4M**), a finding congruent with knockdown of Rab10.

Given the hypothesis that Rab10 and Rab25 function in coordination to deliver LH3 to CIVCs, we would expect to find higher co-localization between the two Rabs when stimulated for secretion. Overexpression of Rab10 WT and Rab25 WT revealed a 50% co-localization in SS media, while only about 30% of the puncta show co-localization in VEGF supplemented media (**Figure 4N and S4A**). Taking a more directed approach, we co-expressed combinations of WT, CA, and DN versions of both Rab10 and Rab25 to further investigate if their co-localization is dependent on activation. Our results demonstrate that expression of the DN Rab10 or Rab25 significantly diminished colocalization compared with any combination of WT or CA, indicating an activation dependency (**Figure 4O and S4B**). To determine if this Rab10 affected LH3 trafficking *in vivo* we compared retina staining between WT and Rab10^+/-^ P6 mice and observed a lack of CIVC co-localization with LH3 **(Figure 4P and 4Q)**.

CIVC vesicles staged for secretion are present as large Col IV puncta that are easily distinguishable from Col IV that is resident in the ER or extracellular environment. Knockdown of Rab10, Rab25 or LH3 (as a negative control) showed a significant reduction in the number of ECs with detectable CIVC vesicles, indicating the loss of Rab10 or Rab25 affects the formation of these structures (**Figure S4C and S4D**). Previous reports determined that vacuolar protein sorting (vps) protein VPS33B was necessary for delivery of LH3 to CIVCs through direct binding of Rab10 and Rab25 [30]. Knockdown of VPS33b significantly reduced Col IV secretion and LH3 trafficking to CIVC vesicles (**Figure S4E-S4H**), consistent with a requirement for Rab10 and Rab25 in trafficking LH3 to CIVC vesicles. Overall, this data suggests that Rab10 and Rab25 work cooperatively to transport LH3 to CIVC vesicles during Col IV secretion (**Figure 4R**).

### Notch signaling regulates LH3 trafficking

Notch signaling in vascular development has been shown to control gene transcription networks critical for blood vessel maturation[19, 31, 32]. Given Col IV BM secretion is associated with more stable, quiescent blood vessels, we sought to determine if Notch signaling intersected with LH3 trafficking via Rab10 and Rab25 to control Col IV secretion. We previously determined that VEGF ligand stimulation largely inhibited Col IV secretion, while SS media greatly increased Col IV secretion (**Figure 3H and I**). In each condition, we assayed for the Notch transcriptional target Hes1 and found that in the SS condition this transcript was significantly elevated, reflecting high-Notch activation (**Figure 5A and S5A**). Using our basal culture media, which contains a proprietary concentration of VEGF, as a control, we compared Col IV secretion in the elevated Notch SS condition to SS media supplemented with either VEGF, or Notch inhibitor DAPT. Serum-starved ECs significantly increased Col IV secretion in reference to the basal media as we previously showed (**Figure 3H**,**I**); however, both VEGF or DAPT administration completely abolished Col IV secretion (**Figure 5B, 5C, S5B, and S5C**). These results demonstrate that Notch signaling is required for Col IV secretion.

**Figure 5.**
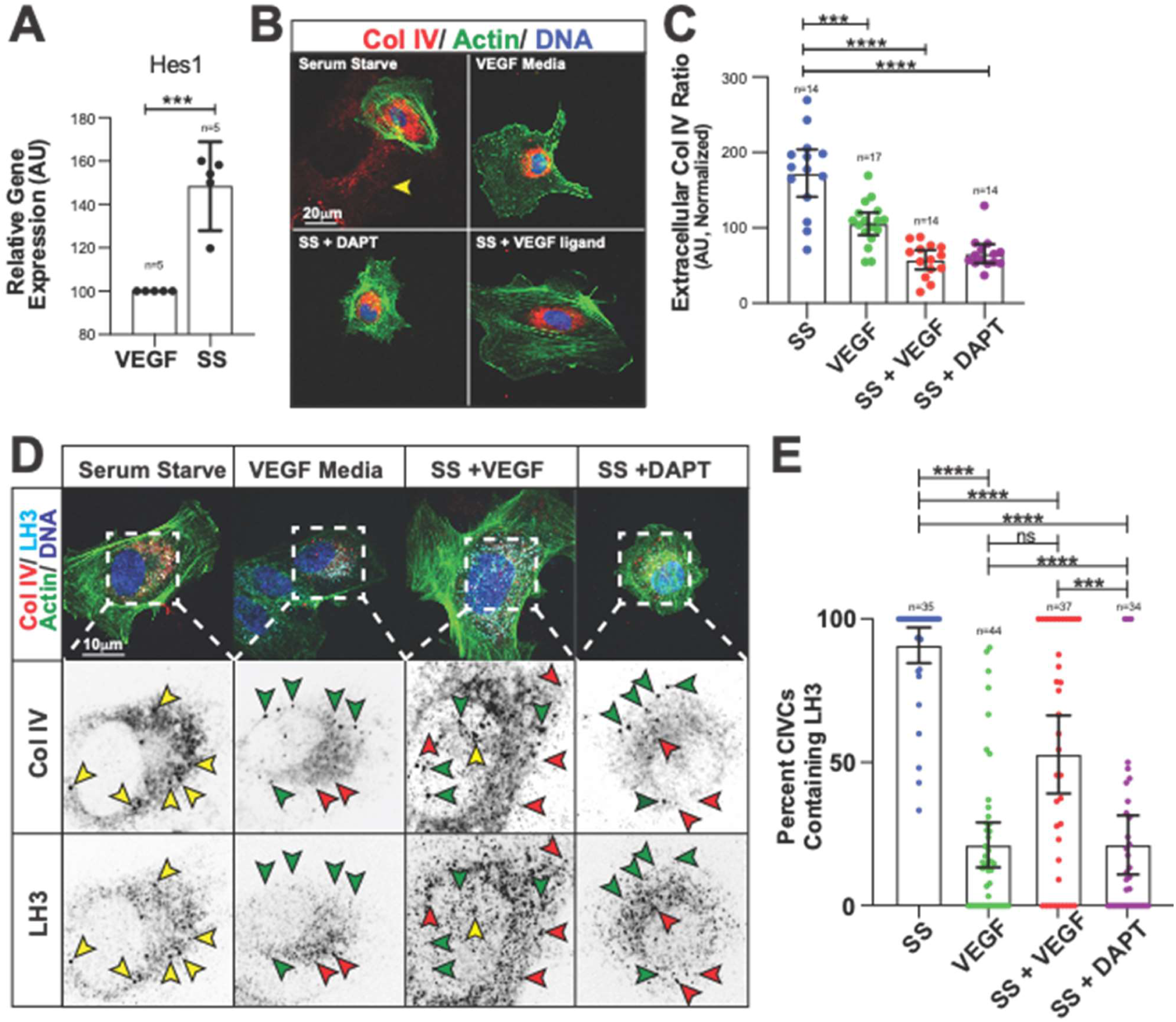
Notch signaling regulated LH3 trafficking. (A) Graph of relative *Hes1* gene expression in ECs cultured in VEGF-containing or SS media. Gene expression levels normalized to GAPDH. (B) Representative images of ECs cultured in VEGF-containing, SS media, SS media + VEGF ligand, or SS media + DAPT and stained for Col IV (red), actin (green), and DNA (blue). Arrowhead denotes extracellular Col IV secretion. (C) Graph of extracellular Col IV ratio of ECs cultured in VEGF-containing, SS media, SS media + VEGF ligand, or SS media + DAPT. (D) Representative images of ECs cultured in VEGF-containing, SS media, SS media + VEGF ligand, or SS media + DAPT and stained for Col IV (red), LH3 (light blue), actin (green), and DNA (blue). Yellow arrowheads indicate co-localized puncta only, green arrowheads indicate Col IV only puncta, and red arrowheads indicate LH3 puncta only. (E) Graph of percent CIVC vesicles co-localized with LH3 in VEGF-containing, SS media, SS media + VEGF ligand, or SS media + DAPT conditions. For all experiments, data are represented as mean ± 95% confidence intervals. Black bars indicate comparison groups with indicated p-values. All p-values are from two-tailed Student’s t-test from at least three experiments. *p≤0.05; **p≤0.01; ***p≤0.001; ****p≤0.0001; ns, not significant.

One potential reason for the lack of LH3 and CIVC vesicle co-localization could be due to a reduction in Col IV expression. To explore this, we cultured ECs in SS media and blocked Notch activation with either DAPT or by adding VEGF and then monitored Col IV transcriptional levels. DAPT and VEGF did not alter transcription of Col IV compared with SS control (**Figure S5D and S5E**). To determine if Notch activity was influencing LH3 trafficking we measured co-localization of LH3 and CIVC vesicles with and without DAPT. Across all conditions, Col IV puncta were present indicating that Col IV transcription was not changed despite Notch inhibition. Notch inhibition dramatically reduced LH3/CIVC vesicle co-localization to an even greater extent than VEGF treatment, in reference to a SS control (**Figure 5D and 5E**). Overall, these data suggest that Notch signaling is required for LH3 transport to CIVC vesicles and downstream Col IV secretion.

### Notch signaling regulates Rab10 GTPase activity through DENND4C

Rabs generally operate in a cascade mechanism where guanine exchange factors (GEFs) convert Rabs from GDP-‘off’ to GTP-bound ‘on’ states [33, 34]. DENND4A,B, and C have been implicated in the activation of Rab10[5, 35]. Interestingly, Rab25 is an atypical Rab that does not have an identified GEF and likely functions more akin to a Rab effector [36, 37]. Our data suggested that activation of Rab10 is required for LH3 transport to CIVC vesicles (**Figure S6A**), thus we explored the idea that Notch governs Rab10 activity through differential GEF expression. First, we surveyed the DENND4 loci for RBPJ Notch-responsive elements in their 5’ cis region (**Figure 6A and 6B**)[38, 39]. Sequence analysis of all three DENND4s revealed that all variants harbored RBPJ binding sites within their respective 0, -2000 5’ untranslated region; however, only DENND4C exhibited a clustering of RBPJ sites close (0, - 500) to the transcriptional start site (**Figure 6B**). Expression analysis between the Notch-low and Notch-high media conditions demonstrated that only DENND4C was significantly upregulated in the SS state (**Figure 6C**), suggesting that Notch activation can directly modulate this transcript. Knockdown of both DENND4A and DENND4C, but not DENND4B, reduced Col IV secretion compared with controls (**Figure 6D and 6E**). However, only DENND4C significantly reduced LH3 co-localization with CIVC vesicles, indicating that DENND4C is likely the major Rab10 GEF required for LH3 trafficking in ECs (**Figure 6F and 6G**). These results indicate that DENND4C is a Notch target. Additionally, this provides further evidence that Rab10 activation by DENND4C is required for LH3 trafficking and Col IV secretion.

**Figure 6.**
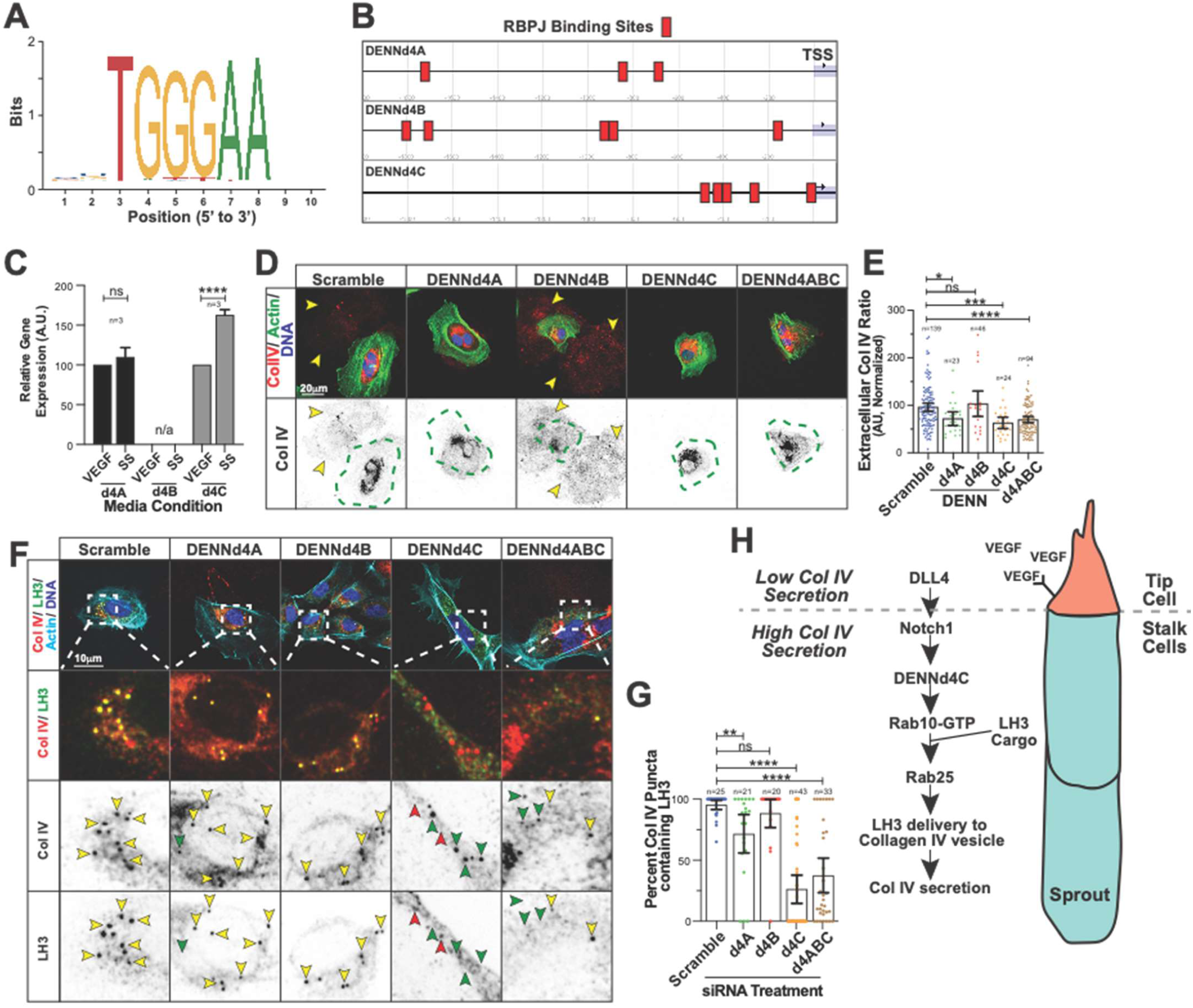
Notch signaling regulated Rab10 GTPase activity though DENNd4C. (A) Predicted RBPJ binding site sequences, identified by the Transfac CSL consensus matrix. (B) Schematic showing RBPJ binding sites upstream of DENND4A (top), DENND4B (middle), and DENND4C (bottom) genes. TSS, transcription start site. (C) Graph of relative gene expression of *dennd4A, dennd4B, dennd4C* in ECs cultured in VEGF-containing or SS media. Gene expression levels normalized to GAPDH. (D) Representative images of scramble, DENND4A, DENND4B, DENND4C, or triplicate siRNA treated ECs stained for Col IV (red), actin (green), and DNA (blue). Dotted green line indicates cell outline. Arrowheads denote extracellular Col IV secretion. (E) Graph of extracellular Col IV ratio in scramble, DENND4A, DENND4B, DENND4C, or triplicate siRNA-treated ECs. (F) Representative images of scramble, DENND4A, DENND4B, DENND4C, or triplicate siRNA-treated ECs stained for Col IV (red), LH3 (green), actin (light blue), and DNA (blue). Yellow arrowheads indicate co-localized puncta only, green arrowheads indicate Col IV only puncta, and red arrowheads indicate LH3 puncta only. (G) Graph of percent CIVCs containing LH3 in scramble, DENND4A, DENND4B, DENND4C, or triplicate siRNA-treated ECs. (H) Schematic diagram of how VEGF signals the tip cell leads to a series of signaling events resulting in the secretion of Col IV from stalk cells. For all experiments, data are represented as mean ± 95% confidence intervals. Black bars indicate comparison groups with indicated p-values. All p-values are from two-tailed Student’s t-test from at least three experiments. *p≤0.05; **p≤0.01; ***p≤0.001; ****p≤0.0001; ns, not significant.

## DISCUSSION

Despite the major biological requirement of Col IV BM for blood vessel integrity and homeostasis, very little is understood about its regulation. Moreover, how Col IV, or other critical BM proteins are regulated by non-transcriptional programs in ECs is largely unknown. To our knowledge, this is the first investigation posing a direct link between the regulation of Col IV secretion in angiogenesis through modulation of trafficking mediators by way of permissive Notch signaling. Our results demonstrate that Rab10 works in combination with Rab25 to transport LH3 to CIVC vesicles staged for secretion. In the absence of Rab10 or Rab25, LH3 transport is halted and Col IV secretion in ECs is dramatically attenuated. Putting this trafficking paradigm into a larger angiogenic framework, we discovered that Notch signaling is required for Rab10 activation, which seems to be the signaling bottleneck for LH3 trafficking and subsequent Col IV secretion. Overall our data illustrate how Notch-based maturation signaling can influence trafficking mediators, providing a critical level of regulation in Col IV secretion during blood vessel development (**Figure 6H**).

Col IV is highly conserved and can be traced down to the earliest bilaterians[40]. Col IV itself is expressed, to some extent, in every vertebrate tissue as an integral BM protein. It is well-established that Col IV is highly enriched in blood vessels and contributes to the overall vascular BM[41]. Col IV is not required for angiogenesis; however, Col IV is required for blood vessel maturation and homeostasis[15]. This enrichment is related to the ability of blood vessels to resist the mechanical strain of circulation. Mutations in Col IV (alpha 1 or alpha 2) in human patients confirm the foundational requirement of Col IV in blood vessel integrity, as the primary clinical manifestation of individuals with SVD is intracerebral hemorrhage. Moreover, SVD patients, who harbor Col IV mutations, demonstrate compromised vessel integrity leading to microbleeds or intracerebral hemorrhages[16]. Rodent experimental models of Col IV mutations strongly echo results in human cohorts in demonstrating that loss of Col IV function or availability results in either embryonic lethality or early postnatal death by way of hemorrhage[15]. These observations clearly indicate that Col IV bioavailability is paramount for blood vessel homeostasis and normal life expectancy.

We investigated post-transcriptional factors that regulate Col IV secretion in ECs and discovered that Rab10 and Rab25 are major trafficking mediators. Rab10 in particular has been implicated in a myriad of processes ranging from GLUT trafficking to regulating ER dynamics[42, 43]. In our hands, Rab10 echoed previous reports in *Drosophila melanogaster* egg chamber development by affecting Col IV secretion; although, our data indicated that Rab10 is an indirect mediator of Col IV trafficking[5]. Our results were congruent with a second report observing that LH3 post-Golgi sorting is controlled, in part, by Rab10 and Rab25 in mouse epithelial cells[30]. Extending these observations in primary ECs, we found that LH3 was indeed required for Col IV secretion, and LH3s delivery to CIVC vesicles was paramount to this process. Interestingly, we also found that Rab10 was the major regulatory step as compared with Rab25. Rab25 is an atypical Rab GTPase that does not possess a canonical GEF for its activation, and thus may behave more akin to a Rab10 effector. While Rab25 was necessary for LH3 trafficking and Col IV secretion, its precise role in this cascade has yet to be determined. On the other hand, Rab10 has three previously reported activating GEFs, DENND4 A, B, and C, requiring a more classical GEF-dependent activation.

Given the stark dependence on Rab10 for LH3 trafficking to CIVC vesicles, we were very intrigued by what upstream mechanisms control Rab10 activity and how they might interface with blood vessel maturation programs. Notch signaling is fundamental to angiogenesis and adult blood vessel homeostasis[19]. In aggregate, Notch activation is repressive, decreasing EC migration and proliferation programs and is generally associated with heightened vessel maturity [19]. In the absence of Notch, blood vessels demonstrate a chronic sprouting phenotype marked by unchecked proliferation and overgrowth[31, 44]. We showed that Notch activation is capable of orchestrating LH3 trafficking to CIVC vesicles by controlling transcription of the Rab10 GEF, DENND4C. This finding has important implications in providing evidence that permissive Notch signaling can also interface with trafficking mediators in a comprehensive top-down regulatory response during angiogenesis.

We observed that the administration of the powerful angiocrine factor, VEGF, effectively shut down Col IV secretion by inhibiting LH3 trafficking via reduction of Rab10 and Rab25 activation. In the context of early blood vessel development, EC migration through tissue is reliant on secretion of BM degrading enzymes, such as MMP9[45]. Energetically, it may be more advantageous to partition ECM breakdown signaling from ECM synthesis signaling as to not mutually undermine each process. In this study, the division between Col IV secretion and degradation was controlled by LH3 trafficking and upstream activation of Notch signaling. It is well established that in a growing sprouts, the leading tip cell has low Notch and the trailing stalk cells have high Notch activation[46, 47]. Our results very closely adhere to this model. In a low Notch state, the tip cell has elevated VEGFR2 expression, thus is experiencing more VEGF signaling and, according to our findings, would likely not be secreting Col IV; potentially shunting more energetic resources to migration and ECM degradation. However, in the stalk cells where Notch activation is elevated, these ECs are buttressing the newly made vascular tunnel by secreting Col IV by, in part, activated LH3 trafficking. To our knowledge, this the first link between Notch signaling and regulation of vascular BM secretion at the post-Golgi trafficking level.

Our results bring into question how Notch may directly impact the trafficking regulation of other critical BM proteins necessary for blood vessel integrity. For instance, others have reported that laminin-111 binds receptors and activates Dll4 signaling[48, 49]. One may speculate that the impact of Notch on the secretion machinery may elicit a feed-forward mechanism in which secretion of BM components binds integrin receptors that reinforce Notch activation. This type of cascade could explain cell-cell independent Notch signaling programs for sustained activation of blood vessel maturity programs required for adult vascular homeostasis. However, the organotypic trafficking programs that govern secretion of Col IV and other BM components are largely un-mapped and will require future investigations.

## ACKNOWLEDGEMENTS

Work was supported by funding from the National Heart Lung Blood Institute (Grant 1R56HL148450-01, R00HL124311) (S.J.G, A.M.W, and E.J.K). This work was also supported by the American Heart Association grant (#18PRE33990097)(S.J.G). We also thank Jennifer Bourne and the Electron Microscopy Center at the University of Colorado Anschutz Medical Campus for assistance with transmission electron micrograph collection and Histotox Labs for tissue processing and staining. Additionally, we thank the Kushner Lab for comments and technical support.

## CONTRIBUTIONS

S.J.G., A.M.W, A.D.P, and E.J.K created zebrafish and cell line constructs. S.J.G. and E.J.K conceived all experiments. R.J. performed mouse retinal experiments. J.R.D. performed histopathology evaluation. S.J.G. and E.J.K wrote the manuscript.

## Material and Methods

### DNA Cloning

Unless otherwise stated, all middle entry vectors were generated by PCR amplification of the desired middle element using attL1/L2-flanked oligonucleotide primers, followed by an LR reaction with either pLenti_705 (17392, Addgene) or pLEX_307 (41392, Addgene). A Gibson assembly was performed with the desired middle elements to be assembled into an EcoRI-BamHI linearized pME-MCS destination. To generate the Rab10 clones, full-length human cDNA Rab10 was synthesized using gene blocks (Operon) and cloned into pME-MCS vector using the primer sequences described in supplementary table 1. Full-length human Rab25 cDNA was purchased from Origene (RC203413, ORIgene) and cloned into pME-MCS as described supplementary table 1. Point mutations were introduced via a Q5 site-directed mutagenesis kit (E0554S, NEB) using primers described in supplementary table 1. All constructs were verified with sequencing.

### Cells and Cell Culture

Primary human umbilical vein cells (HUVECs; PromoCell) were cultured in EBM-2 medium supplemented with 5% fetal bovine serum (FBA), 1% penicillin/streptomycin and 1% growth supplemental kit (EGM-2). Only cells in passages 2-10 were used for our experiments. Human aortic endothelial cells (HAECs) (ACBRI375, Cell-Systems), human brain microvasculature endothelial cells (HBMECs) (ACBRI376, Cell-Systems) and human dermal microvasculature endothelial cells (HDMECs) (CSC2M1, Cell-Systems) were all cultured in EGM-2. The serum starve medium is composed of Optimem (11058021, ThermoSci) supplemented with 1% FBS (25-514, GeneseeSci) and 1% penicillin/streptomycin (P4333, Sigma). Human lung fibroblasts (NHLFs) (CC-2512, Lonza) were cultured in Dulbecco’s modified Eagle’s medium (DMEM) (25-501B, GeneseeSci) media supplemented with 10% FBS and 1% penicillin/streptomycin. Human embryonic kidney cells (HEKs) (85120602, Sigma) were cultured in DMEM media supplemented with 10% FBS and 1% penicillin/streptomycin. All cells were grown at 37C in a humidified atmosphere with 5% CO_2_.

### Zebrafish

Zebrafish (*Danio rerio*) were bred and housed in standard conditions in accordance with the University of Denver. The *Tg(kdrl:eGFP)* (kind gift Victoria Bautch), *Tg(cdh5:gal4FF)* (kind gift Arndt Siekmann), *Tg(5xUAS:tRFP Rab10)* (this study), *Tg(5xUAS:tRFP Rab10*^*Q68L*^) (this study), *Tg(5xUAS:tRFP Rab10*^*T23N*^) (this study). Procedures used in the experiments were approved by the Institutional Animal Care and Use Committee. Morpholinos were purchased from GeneTools LLC and injected using standard protocols. For cryosectioning, zebrafish were fixed in 4% PFA overnight and dehydrated in 100% methanol for 48 hours. Thereafter, embryos were briefly rehydrated in TBST and then incubated in a 30% sucrose solution for 24 hours. Fish were embedded in OCT prior to sectioning and staining as previously reported[50]. Images were obtained using a Leica m165 FC Stereoscope. Zebrafish subjected to TEM were fixed in 4% PFA and 2% glutaraldehyde for 24 hours. TEM processing and imaging were done at University of Colorado TEM core facility.

### Mice

Mice were bred and housed in standardized conditions in the Mouse Research Animal Facility at University of Denver and monitored regularly to maintain a pathogen-free environment. Procedures used in the experiments were approved by the Institutional Animal Care and Use Committee. Rab10^em1(IMPC)J^ mice were obtained from The Jackson Laboratory (MMRRC# 42330). Rab10^em1(IMPC)J^ pups were obtained via intercrossing of heterozygous mutants or via outcrossing with BL6 background mice. None of the intercrossed heterozygote mutant offspring were found to be homozygous null, consistent with other reports[23]. At the time of sacrifice genotypes were determined via tail-clips and PCR.

### Statistics

All statistical analyses were conducted using GraphPad PRISM software. Student’s *t*-test were used to compare the difference between the control and treated group in our studies. A two-tailed P<0.05 was significant, and the data are presented as mean ± 95% confidence interval.

## Supplemental Methods

### DNA Cloning

For co-expression of Rab10 and Rab25 in HUVECs, the destination plasmid pShuttle-CMV (16403, Addgene) was used. To create relative similar levels of expression the two genes of interest were fused together via a p2a viral DNA element. The primers used to clone tRFP-Rab10 and BFP-Rab25 are shown in supplementary table 1. A Gibson assembly was used to assemble all desired elements into the XhoI, EcoRV linearized pShuttle-CMV. All constructs were verified by sequencing.

### Collagen IV Extracellular Secretion Assay

Extracellular Col IV ratio is quantified by taking the total fluorescence intensity of Col IV and the fluorescence intensity inside the cell perimeter. Col IV extracellular ratio = [(total fluorescence – inside fluorescence)/inside fluorescence] *100. A value at 1 or below is representative of little/ no Col IV secretion while a value above 1 is representative of substantial Col IV secretion. Values are then normalized to control condition values. Brefeldin A (00-498093, ThermoSci), chloroquine (C6628, Sigma), cycloheximide (C7698, Sigma) and VEGF (V7259, Sigma) were used for indicated experiments.

### Endothelial Cell Transfection Assay

Cells were transfected using the Neon Transfection System (MPK5000, ThermoSci) according to manufacturer s protocol. Briefly, cells were trypsinized and washed with DPBS then suspended in a solution of R-buffer (100µl; Invitrogen) containing either (100µM) siRNA or (1µg) over-expression plasmids (pLenti_705, pLEX_307 or pShuttle-CMV) using the recommended electroporation protocol (1350 V, 30 ms, 1 pulse). Then, cells were either plated onto pre-treated poly-L-lysine coated glass coverslips for IHC and live-imaging or plated into petri dishes for WB experiments and placed in 37C and 5% CO_2_. IHC and live-imaging experiments were conducted 18-30 hours after transfection while cell lysates were harvested 48-72 hours after transfections.

### Immunohistochemistry

Standard procedures were used for IHC[2]. Briefly, HUVECs grown on poly-L-lysine coated coverslips were washed and subsequently fixed with 4% PFA for 10 min. Cells were then washed and incubated at RT with 0.1% Triton-X for 10 min. Blocking was performed with 2% BSA prior to primary antibody incubation. Commercial antibodies used include: goat anti-Col IV (ab769, Sigma), rabbit anti-Col IV (ab6586, Abcam), rabbit anti-Laminin (L9393, Sigma), mouse anti-heparan sulfate proteoglycan (MABT12, Sigma) and mouse anti-PLOD3 (SAB1400329, Sigma). AlexaFluor conjugated secondary antibodies include donkey anti-goat 555(A32816, ThermoSci) and donkey anti-mouse 647 (A31571, ThermoSci). Hoechst 33342 (H3570, ThermoSci) used as a DNA stain. For wound healing assays, scratches were made when HUVECs were 90% confluent. Dishes were washed twice and then replaced fresh EGM-2 medium for up to 8 hours before fixation with 4% PFA. CellEvent Caspase-3/7 (C10723, ThermoSci) was used to investigate transfection efficiency. All histology was performed at HistoTox Labs (Boulder, CO).

### Sprouting Assay

A fibrin-bead sprouting assay was conducted as described by Nakatsu et al. [22]. Briefly, after trypsinization, HUVECs were incubated with cytodex3 microcarrier beads (C3275, Sigma) at a ratio of 400 cells per bead. The samples were incubated for 4 hours with agitation every 15 min. The mixture was then transferred to a 6cm^2^ dish and cultured at 37C overnight. The next day, beads coated with HUVECs were collected and resuspended in a 2mg/ml fibrinogen solution (F8630, Sigma), which contained 0.15 U/ml aprotinin (A1153, Sigma). As the beads were added to poly-L-lysine pre-treated glass coverslips, 0.625 U/ml thrombin (T4648, Sigma) was added, gently mixed, and incubated at 37C until the gel solidifies. Then, 25,000 NHLFs which were resuspended in 1ml EGM-2 were added on top of the gel. The media was changed every 2 days and fixed 6-8 days after embedding with 4% PFA. Standard IHC staining solutions were used. Images were obtained on inverted Nikon Ti-E spinning disk confocal and analyzed with FIJI software.

### Protein and RNA Isolation from Endothelial Cells

Western blotting was performed using standard procedures. Whole cell lysates were harvested for protein extraction 48-72 hours after transfection. An equal amount (20-35g) of protein was electrophoresed on 12% and 7% polyacrylamide gels and then transferred to nitrocellulose membranes. The membrane was blocked in ∼5% milk or 2% BSA followed by antibody incubation overnight at 4C. Antibodies used are listed below: rabbit anti-α-tubulin (ab52866, Abcam); mouse anti-Rab10 (MABN730, Sigma); rabbit anti-Col IV (ab6586, Abcam). The internal loading control for all experiments was α-tubulin. Secondary HRPs (GeneseeSci) and ProSignal ECL substrate (20-300B, GeneseeSci) were used. For GTP associations, ECs were incubated with indicated media then lysed and incubated with guanosine 5 -triphosphate agarose beads (G9768, Sigma). RNA extraction was performed using TRIzol (15596026, ThermoSci) with standard procedures. RT-PCR was performed on cDNA libraries using high-capacity cDNA reverse transcription kit (4368814, ThermoSci) according to manufacturer instructions. PCRs were performed using ProFlex PCR System (4484073, ThermoSci).

### Generation of Tg(5xUAS:tRFP Rab10) mutant lines

Unless otherwise stated, all middle entry vectors were generated by PCR amplification of the desired middle element using extended and over-hanging oligonucleotide primers, followed by a Gibson assembly with a middle entry plasmid. The tol2 cloning system was used to assemble the p5E 5xUAS promoter and the pME-into a modified 395-destination plasmid (this study).

### Retina Extraction

Eyes from male and female mice were harvested at p6 and fixed in 4% PFA for 2 hours at room temperature. Immediately after fixation, retinas were dissected and flattened by making curve-relieving cuts. The retinas were then fixed for an additional 1-2 hours. Then, retinas were placed in 2% BSA blocking solution overnight at 4C. On day 2, retinas were stained for 24 hours at 4C with goat anti-collagen IV (same as IHC) and rabbit anti-PLOD3 antibody (HPA001236, Sigma). On day 3, retinas were washed twice in TBST and then stained for 24 hours at 4C with conjugated Isolectin B4 (IB4) and Hoechst 33342 (see IHC), donkey anti-goat 555(same as IHC) and donkey anti-mouse 647 (same as IHC). On day 4, the specimens were washed three times in TBST for 10 min and then left in TBST overnight at 4C. On day 5, the retinas were mounted on slides and imaged.

## Supplemental Tables

**Supplementary Table I.**
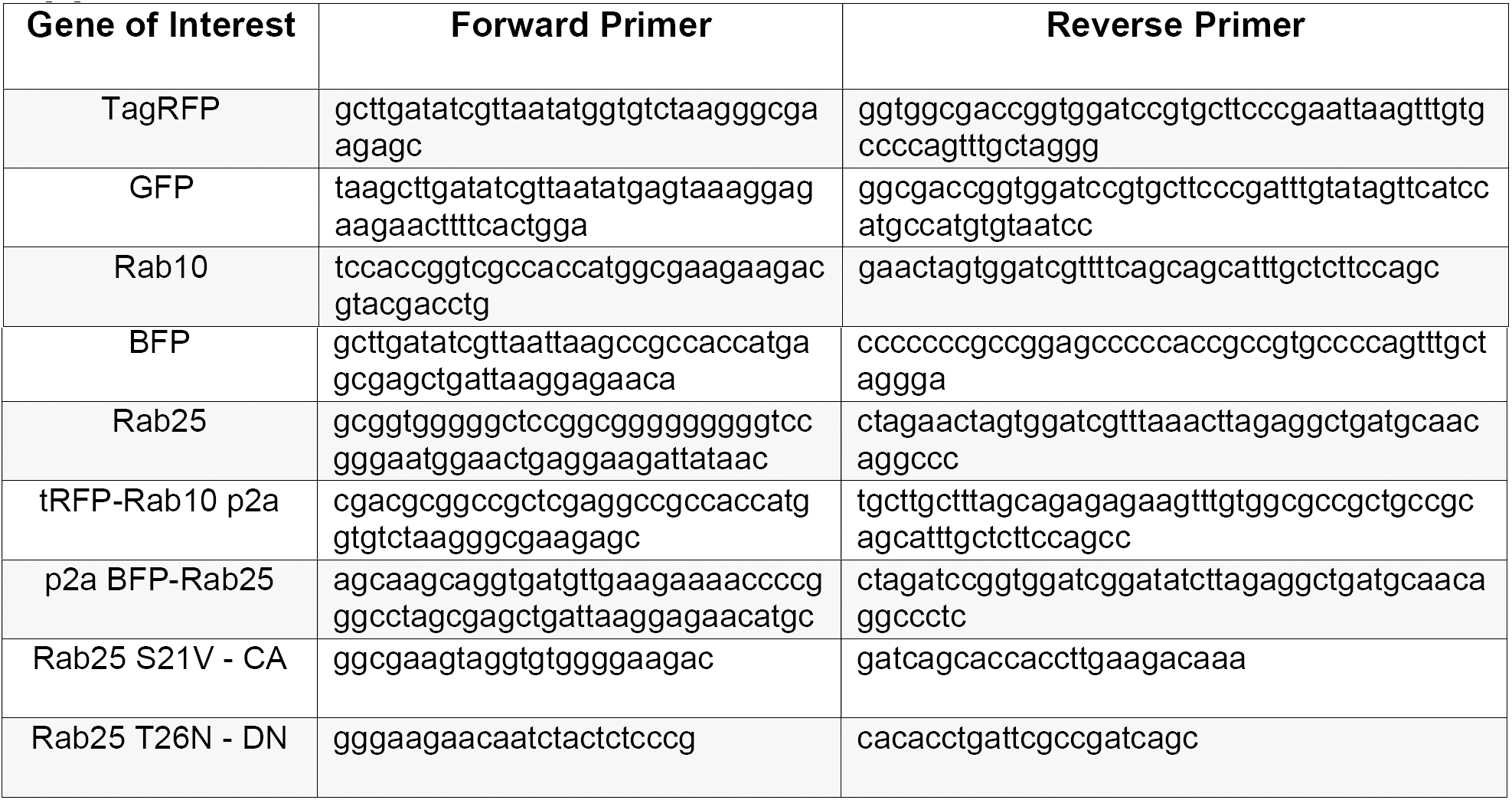
General Cloning Primers. Sequences of the primer pairs used to assemble middle entry plasmids of either GFP or TagRFP versions of Rab10 WT, CA or DN as well as BFP versions of Rab25 WT, CA or DN.

**Supplementary Table II.**
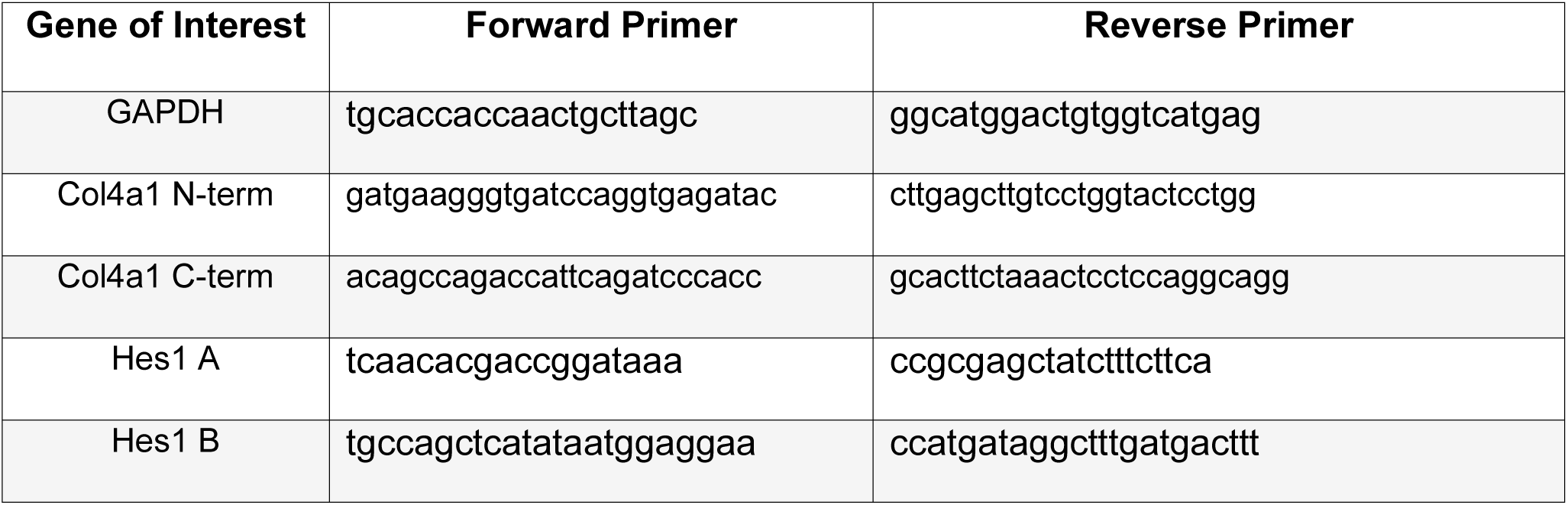
RT-PCR primers. Sequences of the primers used in RT-PCR analysis of gene expression in ECs.

**Supplementary Table III.**
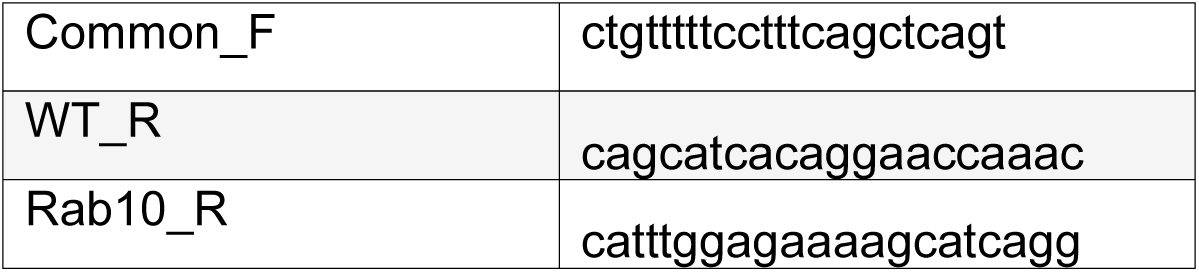
Mouse Genotyping Primers. Sequences of the primers used to determine the genotype of Rab10.

## SUPPLEMENTAL FIGURES

**Supplementary Figure 1.**
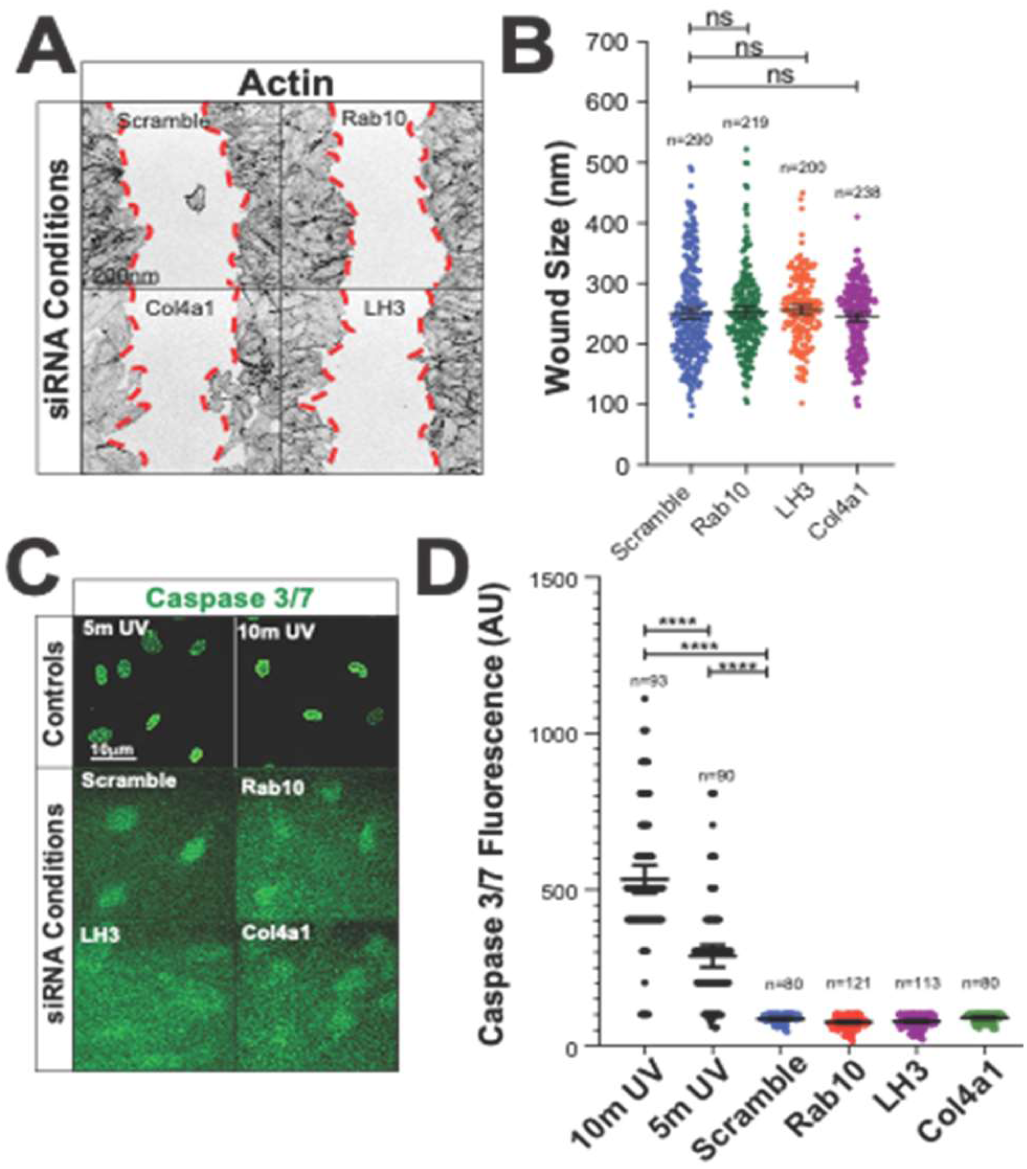
Effect of transfection on EC viability markers. (A) Representative images of scramble, Rab10, lysyl hydroxylase 3 (LH3), and Col IV (col4a1) siRNA treated ECs in scratch wound assay. Cells were stained for actin (grey) to delineate scratch wound margins. Dotted red line indicates wound border. (B) Graph of average wound size in indicated groups of indicated siRNA-treated ECs. n=number of measurements. (C) Representative images of scramble, Rab10, lysyl hydroxylase 3 (LH3), and Col IV (col4a1) siRNA-treated ECs stained for Caspase 3/7 activation (green). Controls were subjected to UV light exposure for indicated times to elicit caspase activation. (D) Graph of Caspase 3/7 activation fluorescence intensity in ECs treated with indicated siRNA treatment groups. Measurement of GFP fluorescence intensity within the nuclei of ECs. n=number of cells. For all experiments, data are represented as mean ± 95% confidence intervals. Black bars indicate comparison groups with indicated p-values. All p-values are from two-tailed Student’s t-test from duplicate experiments. *p≤0.05; **p≤0.01; ***p≤0.001; ****p≤0.0001; ns, not significant.

**Supplementary Figure 2.**
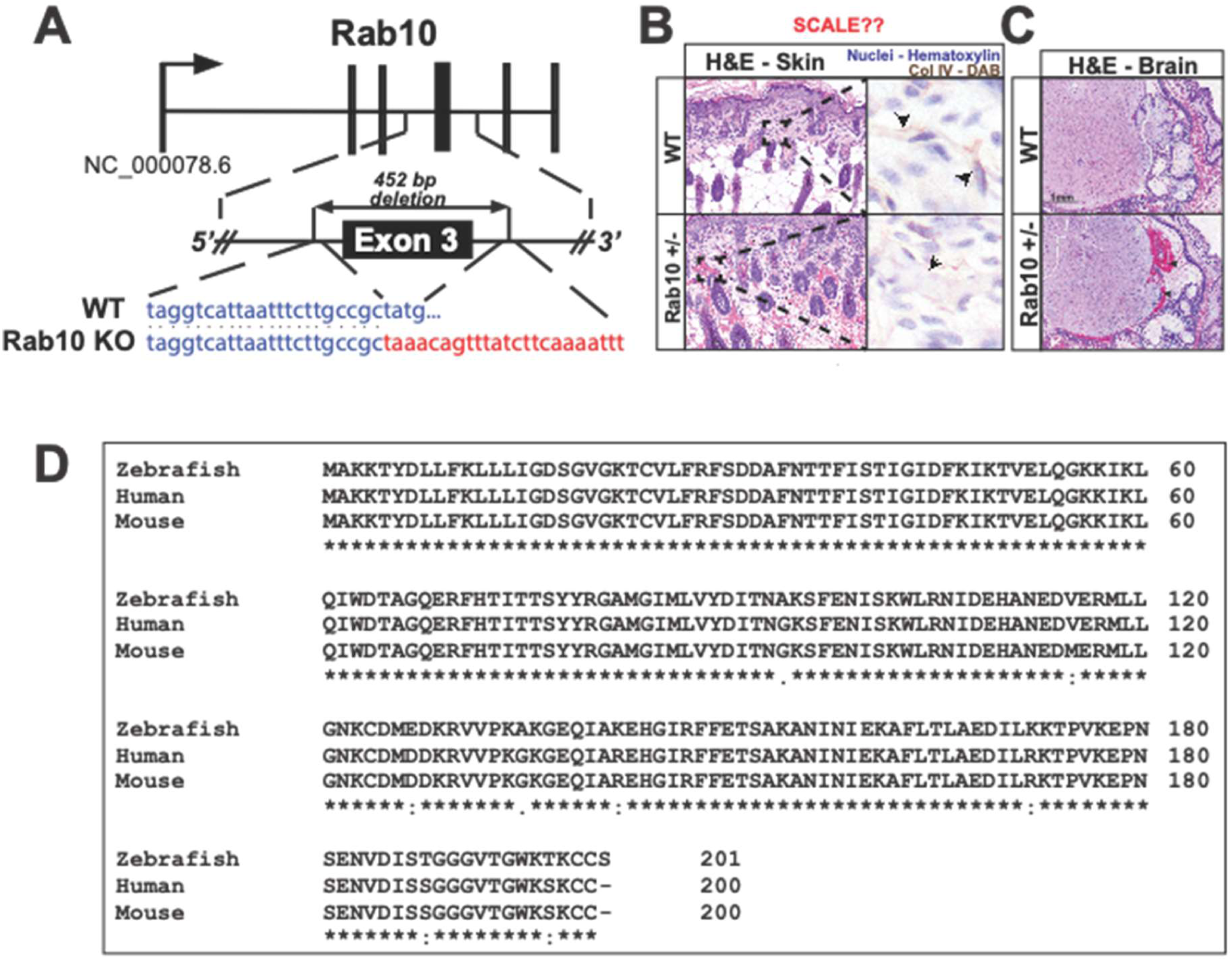
Rab10 influences Col IV bioavailability *in vivo*. SCALE BAR B? (A) Schematic diagram of the CRIPSR mediated *rab10* knockout mouse from Jackson Laboratories. A 452-bp deletion was cloned at exon 3 resulting in an early truncation. (B) Hematoxylin and eosin (H&E) stained skin slices from P6 wild-type or Rab10^+/-^ mice. Collagen IV stained with DAB (3-3’ diaminobenzidine) indicated by arrowheads. (C) Hematoxylin and eosin (H&E) stained brain slices from wild-type or Rab10^+/-^ mice. Arrowheads indicate intracranial/cerebral hemorrhage (bottom). (D) Alignment of *rab10* from human, zebrafish, and mouse.

**Supplementary Figure 3.**
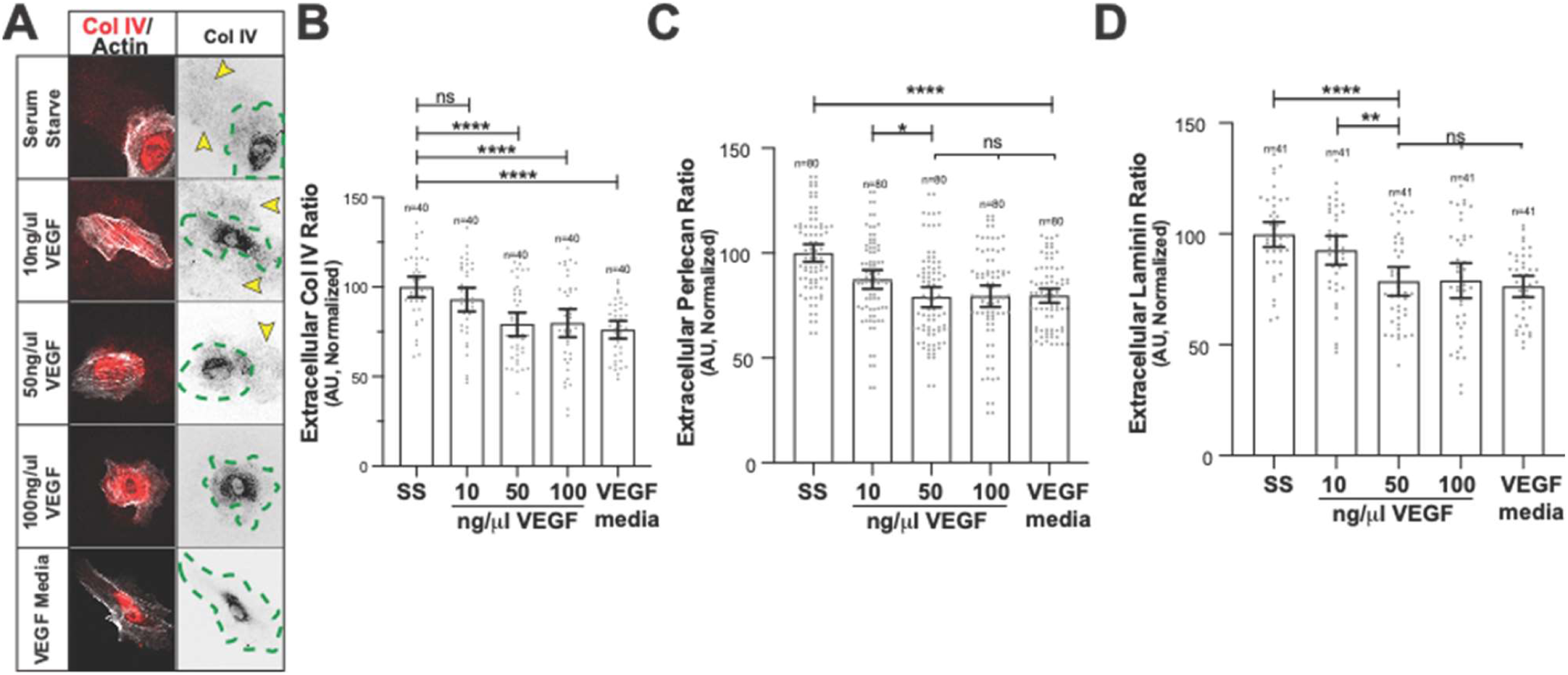
Effect of VEGF on basement membrane secretion in HUVECs. (A) Representative images of ECs cultured in VEGF-containing, SS media, or SS media supplemented with indicated concentrations of VEGF ligand. ECs were stained for collagen IV (red) and actin (grey). Dotted green line indicates cell outline. Arrowheads denote extracellular Col IV secretion. (B) Graph of extracellular Col IV ratio in ECs. (C) Graph of extracellular perlecan ratio in ECs. (D) Graph of extracellular laminin ratio in ECs. For all experiments, data represented as mean ± 95% confidence intervals. Black bars indicate comparison groups with indicated p-values. All p-values are from two-tailed Student’s t-test from at least three experiments. *p≤0.05; **p≤0.01; ***p≤0.001; ****p≤0.0001; ns, not significant.

**Supplementary Figure 4.**
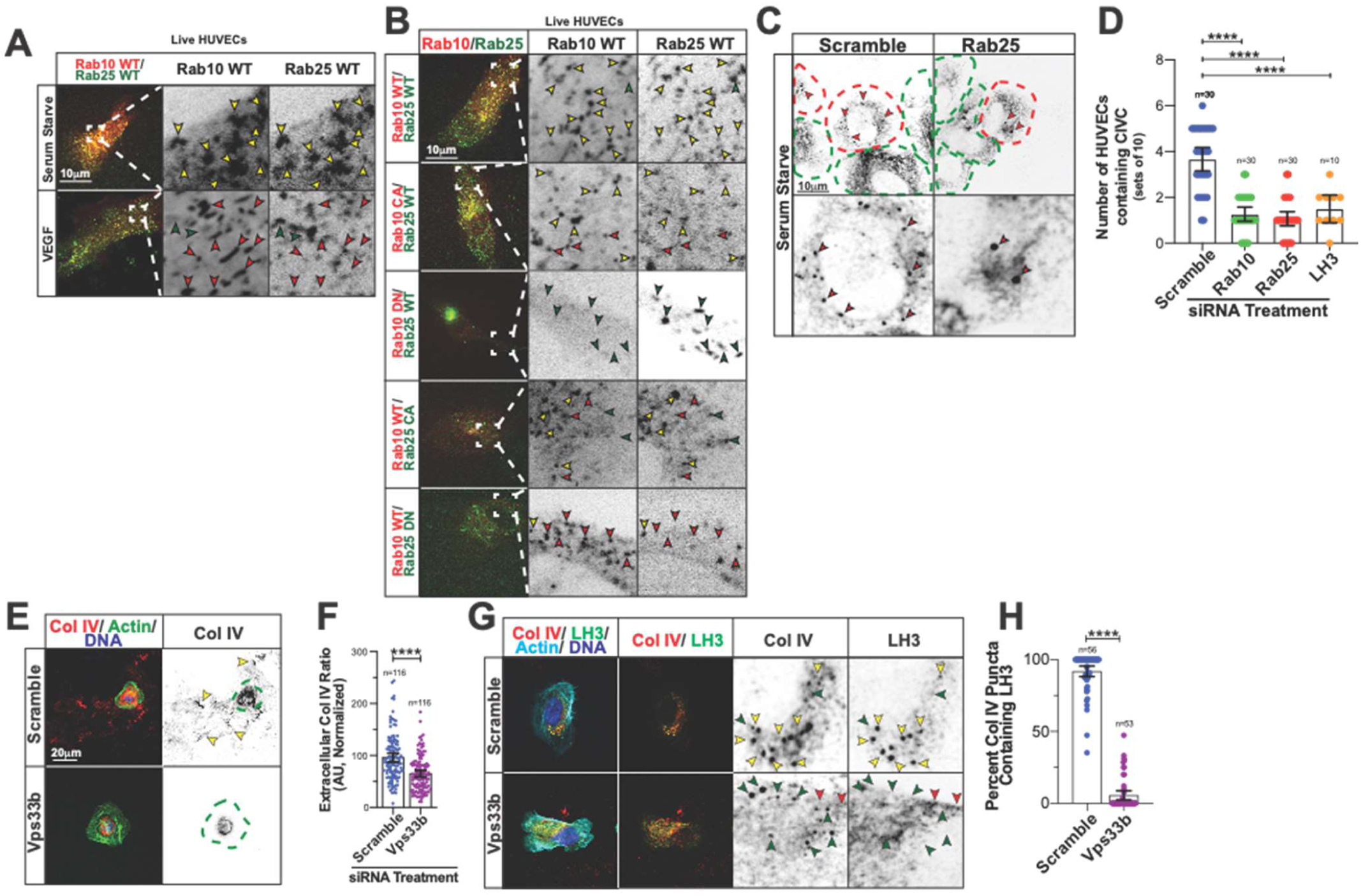
Rab10 and Rab25 work in combination to traffic LH3 to CIVCs. (A) Representative images of ECs co-expressing RFP-Rab10 WT and BFP-Rab25 WT in VEGF containing either VEGF-containing or SS media Yellow arrowheads denote co-localized Rab10 and Rab25 puncta, red arrowheads denote Rab10 puncta only, and green arrowheads denote Rab25 puncta only. (B) Representative images of ECs transfected to co-expression RFP-Rab10 WT, CA or DN and BFP-Rab25 WT, CA or DN cultured in SS media. Yellow arrowheads denote co-localized Rab10 and Rab25 puncta, red arrowheads denote Rab10 puncta only, and green arrowheads denote Rab25 puncta only. (C) Representative images of scramble and Rab25 siRNA-treated ECs cultured in SS media and stained for Col IV (grey). Dotted red and green lines indicates CIVC vesicle positive or negative ECs, respectively. Arrowheads denote CIVC vesicles. (D) Graph of number of ECs containing CIVC vesicles in scramble, Rab10, Rab25, and LH3 siRNA-treated ECs cultured in SS media. (E) Representative images of ECs transfected with either scramble or Vps33b siRNA and stained for Col IV (red), actin (green), and DNA (blue). Dotted green line indicates cell outline. Arrowheads denote extracellular Col IV secretion. (F) Graph of extracellular Col IV ratio of indicated siRNA treated ECs. n, number of measurements. (G) Representative images of ECs transfected with either scramble or Vps33b siRNA and stained for Col IV (red), LH3 (green), actin (light blue), and DNA (blue). Yellow arrowheads indicate co-localized puncta only, green arrowheads indicate Col IV only puncta, and red arrowheads indicate LH3 puncta only. (H) Graph of percent CIVCs containing LH3 in scramble or Vps33b-siRNA treated ECs. For all experiments, data are represented as mean ± 95% confidence intervals. Black bars indicate comparison groups with indicated p-values. All p-values are from two-tailed Student’s t-test from at least three experiments. *p≤0.05; **p≤0.01; ***p≤0.001; ****p≤0.0001; ns, not significant.

**Supplementary Figure 5.**
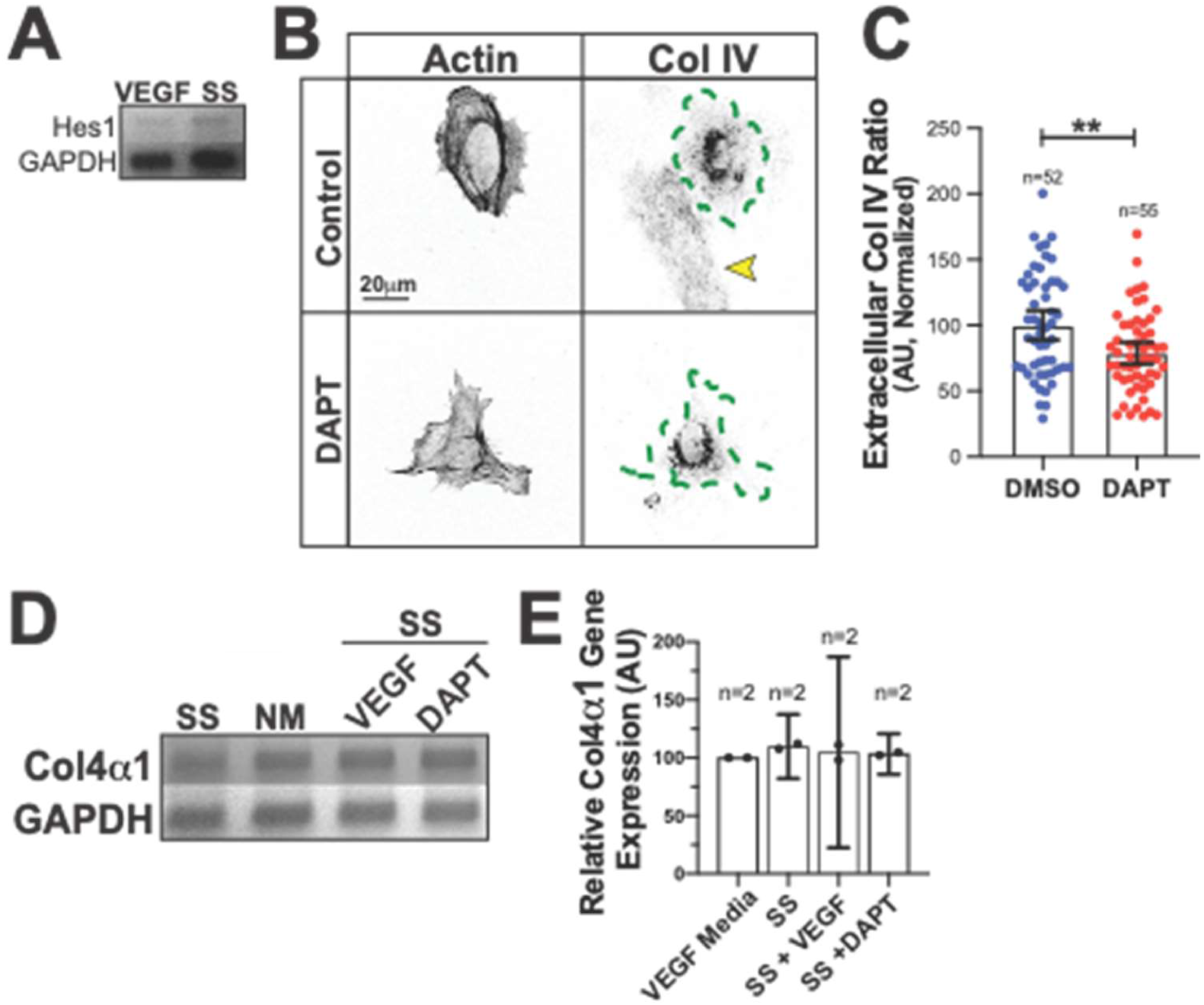
Notch signaling regulated LH3 trafficking. (A) Representative image of *hes1* gene expression ECs cultured in VEGF containing or SS media examined by RT-PCR. Gene expression levels normalized to GAPDH. (B) Representative images of ECs cultured in SS media with either DMSO (control) or DAPT and stained for actin (left) and Col IV (right). Dotted green line indicates cell outline. Arrowheads denote extracellular Col IV secretion. (C) Graph of extracellular Col IV ratio of ECs cultured in SS media with either DMSO (control) or DAPT. (D,E) Representative RT-PCR ad graph of Col4a1 expression between indicated groups. n= number of experiments. For all experiments, data are represented as mean ± 95% confidence intervals. Black bars indicate comparison groups with indicated p-values. All p-values are from two-tailed Student’s t-test from at least three experiments. *p≤0.05; **p≤0.01; ***p≤0.001; ****p≤0.0001; ns, not significant.

**Supplementary Figure 6.**
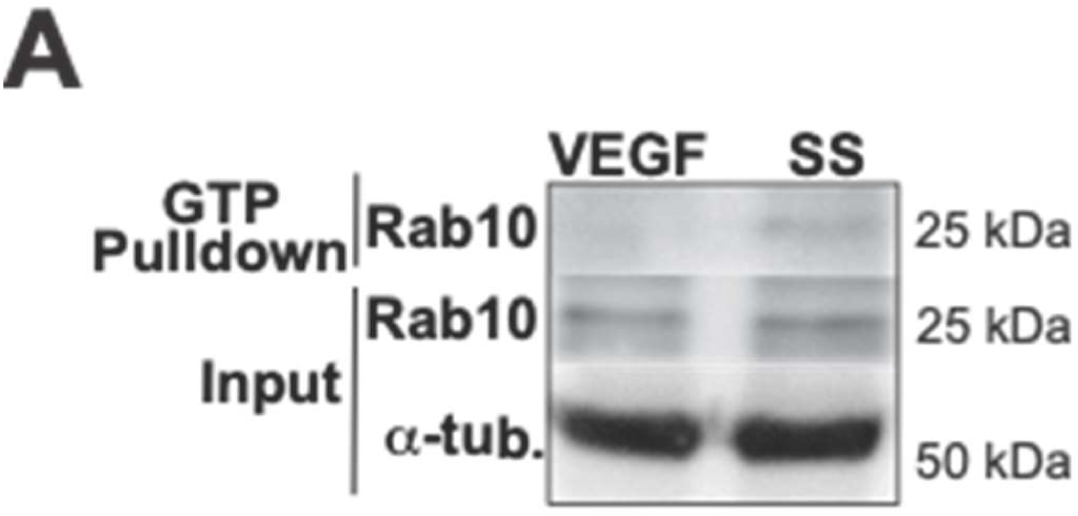
Effect of Vps33b siRNA on Col IV secretion in ECs. (A) Immunoblot of glutathione-bead pulldowns from ECs expressing endogenous Rab10 cultured in either VEGF containing or SS media.

